# A characterization of piARNs, their biogenesis and their targets in *Spodoptera frugiperda* (Lepidoptera, Noctuidae)

**DOI:** 10.1101/2025.09.10.675288

**Authors:** Imène Seninet, Mélanie Gasser, Sylvie Gimenez, Fabrice Legeai, Karine Durand, Nicolas Nègre, Kiwoong Nam, Bernard Duvic, Emmanuelle d’Alençon

## Abstract

PIWI-interacting RNAs (piRNAs) and PIWI proteins were initially described as involved in gametogenesis and preservation of genome integrity through the control of transposable elements (TE). Expressed also in the soma and able to regulate protein coding gene expression, they are involved in multiple biological pathways including host-pathogens interaction, sex determination, reproductive isolation. *Spodoptera frugiperda* is a major invasive insect pest species consisting of two strains with different host-plant ranges. In this paper, we characterized proteins and genomic regions involved in their biogenesis as well as TE and gene transcripts regulated by piRNAs.

By phylogenetic analysis, we identified two new *Piwi* genes conserved in the genus *Spodoptera*, compared to the Lepidopteran model *Bombyx mori*, one more than in *Drosophila*. One of them, more expressed in gonads then soma could be a functional homolog of Drosophila PIWI or replace AGO3. A pool of 11 sRNA-Seq libraries was used to annotate piRNA clusters with Shortstack in the genomes of the two strains of *Spodoptera frugiperda*. Identification of TE targeted by piRNAs revealed that active transposons differ between the two strains of *S. frugiperda* despite of a similar TE content, as putative cause or consequence of reproductive isolation. GO analysis of genes targeted by piRNAs shows that some are involved in protein translation initiation. A piRNA cluster in the *Masc* gene suggests that sex determination is regulated by piRNAs in *Spodoptera frugiperda*.

Our analysis supports that piRNAs have additional roles than silencing of transposable elements and contributes to functional annotation of the two strains genomes of *Spodoptera frugiperda*.

## 1. Introduction

Piwi-interacting RNAs (piRNAs) are a class of small non-coding RNAs known to preserve genome integrity by silencing transposable elements (TE) in the germline of animals (^1–4^ for review). Piwi protein was initially identified as necessary for germ stem cell self-renewal in *Drosophila* ^5,6^. PiRNAs were first described in *Drosophila* involved in the process of Su(Ste) silencing of Stellate ^7^. Since that time, in addition to TE, piRNAs have been shown to regulate cellular genes involved in several developmental pathways in animals (^8,9^ for review).

Their action is not restricted to the germline, they have been reported to function also in somatic tissues for instance in *Drosophila* in the fat body ^10^ and the brain ^11,12^. In addition to repression of TEs, somatic piRNAs are involved in defense against RNA viruses in the mosquito *Aedes aegypti* ^13^. Lewis *et al.* recently found somatic piRNAs targeting TEs and cellular mRNAs in 20 species of Arthropods showing that this somatic pathway is an ancestral trait in their phylogeny ^14^.

Like other small non coding RNAs, piRNAs have a specific size range and interact with proteins of the Argonaute family which they guide to their regulatory targets by homology, typically leading to decrease in their expression. The Argonaute family of proteins consists of two clades, the AGO clade and the PIWI clade. AGO clade proteins complex with microRNAs (miRNAs) and small interfering RNAs (siRNAs), which derive from double-stranded RNA (dsRNA) precursors ^15,16^. Proteins of the PIWI clade interact with piRNAs. The copy-number of the *Piwi* gene family varies among animals suggesting possible sub-functionalisation or neo-functionalisation, for example *C. elegans* encodes two PIWI proteins from a lineage specific duplication ^17^, *Drosophila melanogaster* encodes three PIWI proteins PIWI, AUBERGINE (AUB) and AGO-3 ^14^, while annotation of the *A. pisum* genome identified eight copies of the *Piwi* genes and two copies of the *Api-ago*3 ^18^ and some *turbellaria* flatworms show up to eight lineage specific paralogs ^19^.

The piRNA biogenesis occurs by two pathways, the primary pathways and the ping-pong amplification loop, the first one provides a pool of piRNAs against all tranposable elements (TE) hosted by the animal population, whereas the second allows enrichment of piRNAs against active transposons. piRNAs are produced from long single-stranded RNAs transcribed from genomic loci containing remnants of TE, known as piRNA clusters. These transcripts, usually antisense to TE transcripts, are processed into primary piRNAs by Zuchini endonuclease ^20^. They are loaded into PIWI proteins which cleave sense TE transcripts generating secondary piRNAs, which in turn guide AGO-3 to cleave piRNA cluster transcripts through an amplification loop called the Ping-Pong loop ^21,22^. PIWI-class members exhibit strand specific interaction with piRNAs, PIWI and AUB are associated with antisense TE-piRNAs; while AGO-3 is associated with sense TE-piRNAs ^21,22^. Primary piRNAs have a strong bias for uridine residue at the 5’end ^23,24^, while secondary piRNAs have a strong bias for adenine at position ten (10A), and show complementarity over 10 bp to primary piRNAs ^22^.

*Spodoptera frugiperda* also called Fall armyworm (FAW) is a Lepidopteran pest of crops, described as two strains with different host-plant range ^25,26^, the corn strain mostly found on corn (SfC), the rice strain associated to grasses (SfR). Since publication of their genomes ^27^, miRNAs genes have been annotated ^28,29^ but almost nothing is known about the piRNA pathway except that PIWI protein homologs play a role in antiviral response ^30^. Another study of the DNA or non-retroviral RNA viruses embedded in piRNA clusters of 48 Arthropods mentioned that some Rhabdoviridae are enriched in piRNA clusters in the genome of the ovary cell line *Sf*9 ^31^ without giving any details on the piRNA clusters themselves. In the silkworm, the lepidopteran model, the ping-pong mechanism has been deciphered *in vitro* in cultured *Bm*4 germ-cells ^32–34^. *Bombyx* lacks orthologs of Drosophila PIWI and has no transcriptional silencing machinery that relies on phased piRNAs. It has only AGO3 and AUB which are cytoplasmic and involved in post-transcriptional silencing of TEs or other gene targets ^32^. Among the other gene targets of piRNAs in *Bombyx mori*, the sex determination gene *Masculinizer* is cleaved in females by a piRNA encoded by W chromosome resulting in female developmental process ^35^. In this paper, we present a phylogenetic analysis of the Argonaute proteins in insects and the expression pattern of *Spodoptera frugiperda* genes in soma and gonads. We annotated piRNA clusters in the two strains genomes of *Spodoptera frugiperda* thanks to a dataset of sRNA-Seq from the two strains. We then identified TE regulated by piRNAs in the two strain genomes and compared the patterns obtained. Finally, we focused on the genes of the insect regulated by piRNAs and explored their functional annotation and putative involvement in sex-determination.

## 2. Material and Methods

### 2.1 Biological material

The sRNA-Seq data described in ^36^ and re-used in this study were obtained from two laboratory strains of *S. frugiperda*: the corn-strain, originating from French Guadeloupe and the rice-strain, originating from Florida, USA, both from the native area. Each strain was reared on its principal and alternative host plant (*Zea mays* L. cv B73 or *Oryza sativa* L. japonica cv Arelate) under controlled conditions (temperature: 24°C, photoperiod 16:8 light:dark, relative humidity: 65%). Eggs from *S. frugiperda* were deposited on plant leaves and fed ad libitum. After three generations on plants, larvae of fourth instars (L4) were collected. Therefore, larvae selected for small RNA extraction and sequencing comprised four groups: C strain reared on corn (CC), R strain reared on corn (RC), C strain reared on rice (CR) and R strain reared on rice (RR). Each experiment comprised two independent replicates. To obtain sRNA-Seq from three developmental stages (this study), the corn-strain was reared on the artificial diet “Poitout” ^37^ under the same controlled conditions.

### 2.2 Identification of Sf Argonaute proteins and phylogenetic analysis

To identify Argonaute genes in the genome of *Spodoptera frugiperda* (version 6.0), we used as query the RNA interference genes already annotated in a previous version ^38^ in a search by TBLASN in the SfruDB database http://bipaa.genouest.org/sp/spodoptera_frugiperda/ and annotated the corrresponding transcripts in Webapollo. We corrected the exon/intron structure based on homology. The Argonaute proteins IDs of all insects used in this study are available in Supplementary Table 1. The evolutionary history was inferred using the Neighbor-Joining method ^39^. The evolutionary distances were computed using the Poisson correction method ^40^. Evolutionary analyses were conducted in MEGA11 ^41^.

### 2.3 Analysis of differential gene expression

To study somatic and germinal expression of Argonaute genes, we re-used RNA-Seq data reported in recent studies ^42, 43^. Briefly, for each replicate (three for the fat body, two for the gonads of *S. frugiperda*), total RNA was extracted using the EZNA Total RNA kit I (Omega) from the fat bodies of 25–28 larvae and the testes of five larvae. Library preparation and RNA sequencing (125 bp paired-end reads generated on an Illumina HiSeq 2500 platform) were performed by the Montpellier GenomiX (MGX) sequencing facility (CNRS, Montpellier, France). Raw reads were aligned with the sfC.ver5.fa reference genome sequence for *Spodoptera frugiperda* ^44^ with STAR v20201 according to ^43^. We assessed differences in gene expression between insect tissues (male gonads vs. fat body in fifth instar larvae), using the read count data as input for the R package DESeq2 ^45^. DESeq2 uses negative binomial generalized linear models to test for differential expression. An adjusted *p*-value for multiple testing was computed with the Benjamini-Hochberg procedure to control for false discovery rate (FDR). Results with FDR<0.05 were considered statistically significant.

### 2.4 Small RNA extraction, library preparation and sequencing

In a previous paper, we described microRNAs of the corn and rice stain of *Spodoptera frugiperda* reared on plants, corn and rice ^28^. In the present paper we re-used the same sRNA-Seq dataset to study piRNAs in SfC and SfR. These datasets had been obtained in the following conditions: Briefly, small RNA was extracted from 12 larvae at the 4^th^ instar (two biological replicates) using the mirVana Isolation kit (Ambion). cDNA library preparation was made using the TruSeq ®Small RNA Sample Prep Kit (Illumina) and sequencing on Illumina HiSeq 2500. A size selection in the range of 15 to 40 nt was performed. Library preparation and sequencing (Single reads, 50 bp) were performed by the platform MGX-Montpellier GenomiX (Montpellier, France). To identify piRNAs candidates involved in sex-determination, additional sRNA-Seq data from three developmental stages (eggs, larvae, pupae) of SfC were obtained, following the same protocol except that the sequencing was performed on the GATC Biotech platform on Illumina HiSeq2000.

### 2.5 Size profiling and mapping of sRNAs to various genomic references

Size profiling of sRNA-Seq reads (Trimmed FastQ files) mapping to the reference genome of SfC (version 5.0) or SfR (version 3.0) was performed using the Galaxy pipeline sRNApipe ^46^. Piecharts representing their distribution on various genomic references were obtained with the same pipeline. Briefly sRNApipe aligns sRNA-Seq reads on the reference genome, these reads that align are called “bonafide reads”. The length of these reads are computed and then they are aligned to various genomic references (miRNA precursors, transcripts, TE consensus, rRNA, snoRNA, tRNA) and counted.

### 2.6 Identification of piRNA clusters

Trimmed FastQ files corresponding to smallRNAs extracted from SfC (or from SfR) were combined and filtered to keep only sequences of size range comprised between 24 and 39 nt, expected for piRNAs. These pools were used as input for the software Shortstack version 4.0.3 ^47, 48, 49^ with default settings except -dicermin 24 and -dicermax 39 defining the minimal and maximal lengths of piRNAs. Shortstack is publicly unvailable at https://github.com/MikeAxtell/ShortStack/releases and performs i) alignment of the reads using Bowtie ^50^ against the reference genome SfC version 5.0 (or SfR, version 3.0, respectively) ii) computation of the depth of small RNA coverage along the reference genomes iii) annotation of piRNAs precursors corresponding to overlapping genomic regions where the depth of piRNA coverage is greater or equal to 1 read per millions (mindepth by default) according to ^51^. https://github.com/MikeAxtell/ShortStack/blob/master/README.md#usage

### 2.7 Identification of Long Non-coding RNAs(LNCR)

After trimming, RNA-Seq from hemocytes, fat body ^43^ and gonads ^42^ were aligned against the genome reference of *S. frugiperda* version 5.0 using STAR version v2.5.2a. The BAM files were merged using Samtools. The alignment results were assembled using StringTie version 2.1.4 with stringtie_ignore_gtf parameter. Identification of candidate lncRNAs from the assembled transcriptome was based on applied filters: exon number ≥2, transcript length >200 bp, using the FEELnc pipeline version 0.2 ^52^. Prediction of the protein-coding potential was also evaluated using FEELnc using OGS5.0. We could identify a total of 5755 unique sequences of LNCRNAs. The LNCR homolog to TE consensus were identified by Blastn.

### 2.8 Annotation of transposable elements

The annotation of TE in SfC and SfR genome assemblies (v5.0 and v3.0, respectively) was performed using REPET v2.5 ^53, 54^. Following a Blastn of the genome against itself, the TEdenovo pipeline ^53^ detects high scoring pairs, which are clustered according to various methods before identification of a consensus sequence representative of each type of repetitive element. For the classification of consensus sequences according to ^55^, various features based on homology or structure were taken into account. Consensus sequences were built only if at least 3 similar copies were detected in the genome. In a second step, the TEannot pipeline ^54^ was used to annotate TE in the genomes. The consensus from TEdenovo were used in a first round of annotation of each genome SfC and SfR independently. Then a second round of TEannot was performed using a pool of SfC and SfR TE families with full length consensus fragments. This second round of TE annotation allowed identification of TE specific of one strain or the other from their genome coverage.

### 2.9 Mapping of piARNs on TE consensus and gene models of Spodoptera frugiperda

Trimmed reads of sRNA-Seq were aligned on different reference sequences using Bowtie1 ^50^ version v0.12.7. For mapping on TE consensus 2 mismatches were allowed –n 2. Mapping with perfect match –n 0 were performed on coding or non coding gene transcripts or genomes. On each reference, mapping was performed with reads mapping uniquely –m 1 or with multimappers. The number of reads mapping to various references was counted using Samtools idxstat version v.0.1.18.

### 2.10 GO term enrichment

The gene functions and associated Gene Ontology (GO) terms of piRNA sequences were determined by first aligning piRNA sequences to the coding sequences of *Spodoptera frugiperda* genes from the OGS6.1 genome assembly downloaded from the BIPAA platform using Bowtie 1 ^50^. A strict threshold was applied to identify highly targeted genes (≥100 mapped sequences). Functional enrichment analysis was then performed on these target genes using BLAST2GO v6.0.3 ^56^ with default parameters. We identified overrepresented GO terms using Fisher’s exact test with a False Discovery Rate threshold < 0.1.

### 2.11 Identification of ping-pong signatures

To detect pairs of piRNAs showing a ping pong signature, we used PingPongPro, a script developed at the Institute of Molecular Biology gGmbH in Mainz (Germany). It is freely available at https://github.com/suhrig/pingpongpro/releases. This software uses alignment files, either the sequence Alignment Map (SAM) or the Binary-compresses Sequence Alignment/map format (BAM) of small non coding RNAs on the reference genomes to detect stacks on both strands overlapping by 10 nucleotides, starting with a U at their 5’end and with a A at position ten. Then it calculates a score for the validity of the signatures. For the command line tool, we used BAM alignment files.

## 3. Results

Biogenesis of piRNAs mainly involves Argonaute proteins and precursor RNAs resulting from transcription of piRNAs clusters. We thus started to explore these two components of the pathway.

### 3.1 Phylogenetic analysis of Argonaute proteins in insects

We first identified Argonaute protein genes in the genus *Spodoptera* and analyzed their number and phylogenetic relationships with those of other Lepidopteran species and of other insect orders.

*Ago*1, *Ago*2, *Ago*3 show a 1:1 orthology relationship between *Spodoptera frugipe*rda and *Bombyx mori*, the Lepidopteran model, whereas *Spodoptera frugiperda* has three Aubergine genes instead of one in *Bombyx mori*. These three Aubergine genes *Aub*1, *Aub*2, *Aub*3 are present in all species of the genus *Spodoptera* studied. In their paper on the role of PIWI in antiviral responses, Xia *et al.*, identified additional copies of Aubergine genes in the genome of an invasive population of *Spodoptera frugiperda* ^30,58^. We studied their phylogenetic relationships with the genes identified in the reference genome of native populations (Supplementary Figure 1). They found two proteins clustering with our AUB1 and two others clustering with our AUB3 (only one gene like us for AUB2). We think that these additional copies either evolved during the very recent process of worldwide invasion or that they result from artefactual duplication during the genome assembly process because of high heterozygosity (see Discussion section).

### 3.2 Germinal versus somatic expression of Argonaute proteins

Until 2016, the piRNA pathway was supposed to suppress TE activity in gonads only, when Jones *et al.* showed that it was also functional in somatic tissue *i.e.* the fat body of adult *Drosophila* ^10^. The *Piwi* mutant showed depletion of fat body piRNAs and increased TE mobilization in this tissue too. This led us to explore *Ago* gene expression in gonads and fat body in our insect species. For that purpose, we relied on the RNA-Seq analysis described in a recent paper ^42^. Briefly, we obtained a mean of 23.9 million reads per replicate (2 for gonads, 3 for fat body), about 18.6 million of which (78%) mapped to unique sites on reference genome version 5.0 of *Spodoptera frugiperda* ^42^. The gene expression profiles of the gonads and fat bodies were compared with the DESeq2 software, which can be used to analyze differential gene expression between conditions based on read counts mapping to particular coding sequences, here OGS6.0 from *Sf*C genome version 5.0. We performed principal component analysis (PCA; Supplementary Figure 2A) on the various samples (tissues, replicates). The log2 fold-change for each gene was plotted as a function of mean expression — *i.e.* the mean number of counts normalized by size factors (MA plot; Supplementary Figure 2B). Blue dots correspond to genes displaying significant differential expression between the gonads and the fat body, at a FDR of less than 0.05. The heatmap on Figure 2 shows that genes *Ago*1, 2, 3, and *Aub*1, 2 are expressed in both tissues. *Ago*1 and *Ago*2 are significantly more expressed in the soma than in gonads (Supplementary Table 2, Tab AGO, DESeq2 results). On the contrary, *Aub*1 and *Aub*2 are significantly more expressed in gonads than fat body, whereas *Ago*3 is expressed to the same level in gonads and fat body. *Aub*3 is expressed at a very low level in both tissues.

**Fig. 1.**
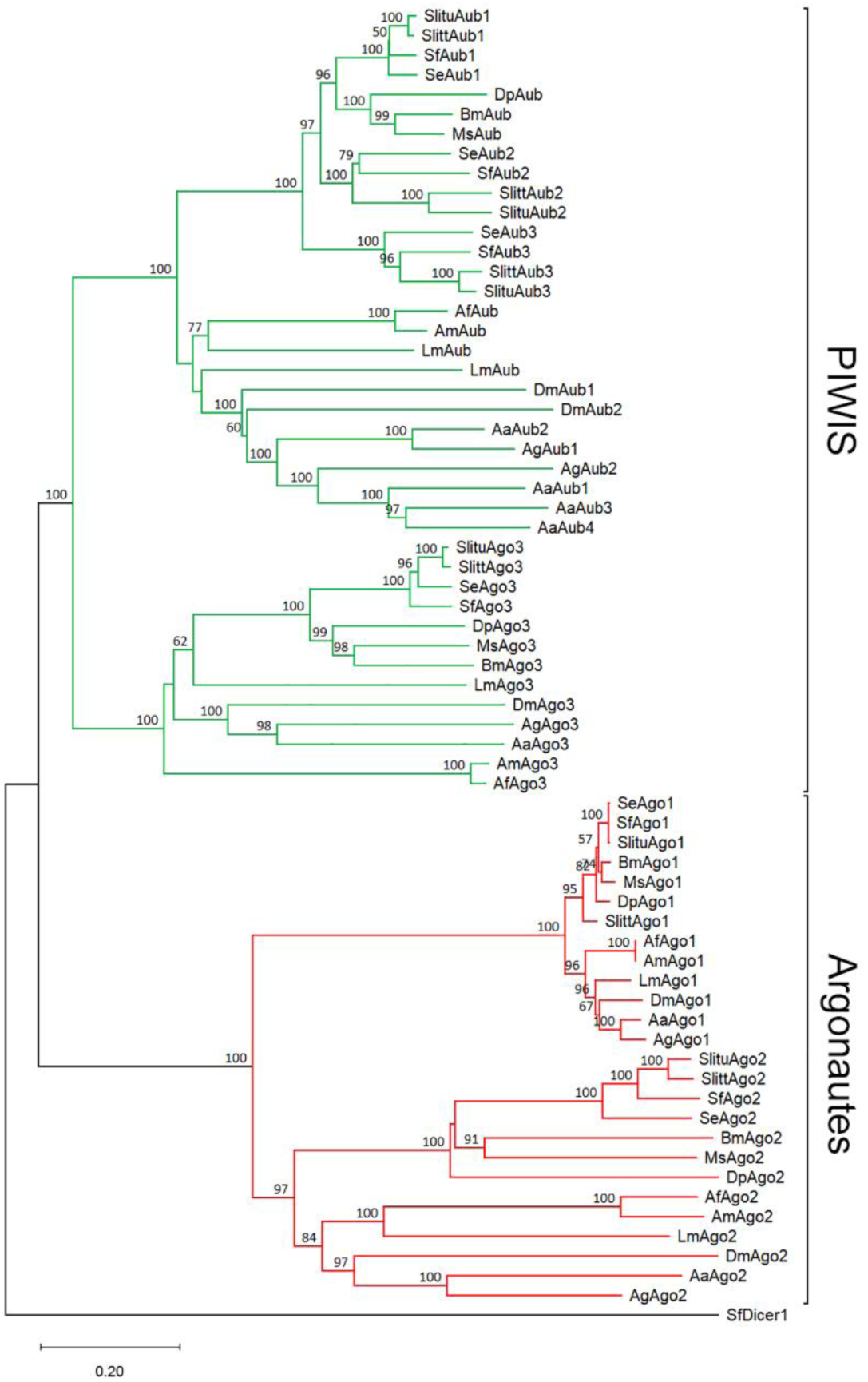
Evolutionary relationships of taxa. The evolutionary history was inferred using the Neighbor-Joining method ^39^. The percentage of replicate trees in which the associated taxa clustered together in the bootstrap test (1000 replicates) are shown above the branches ^57^. The tree is drawn to scale, with branch lengths in the same units as those of the evolutionary distances used to infer the phylogenetic tree. The evolutionary distances were computed using the Poisson correction method ^40^ and are in the units of the number of amino acid substitutions per site. This analysis involved 65 amino acid sequences, whose accession number of sequences are available in Supplementary Table 1. All ambiguous positions were removed for each sequence pair (pairwise deletion option). There were a total of 2697 positions in the final dataset. Evolutionary analyses were conducted in MEGA11 ^41^.

**Figure 2.**
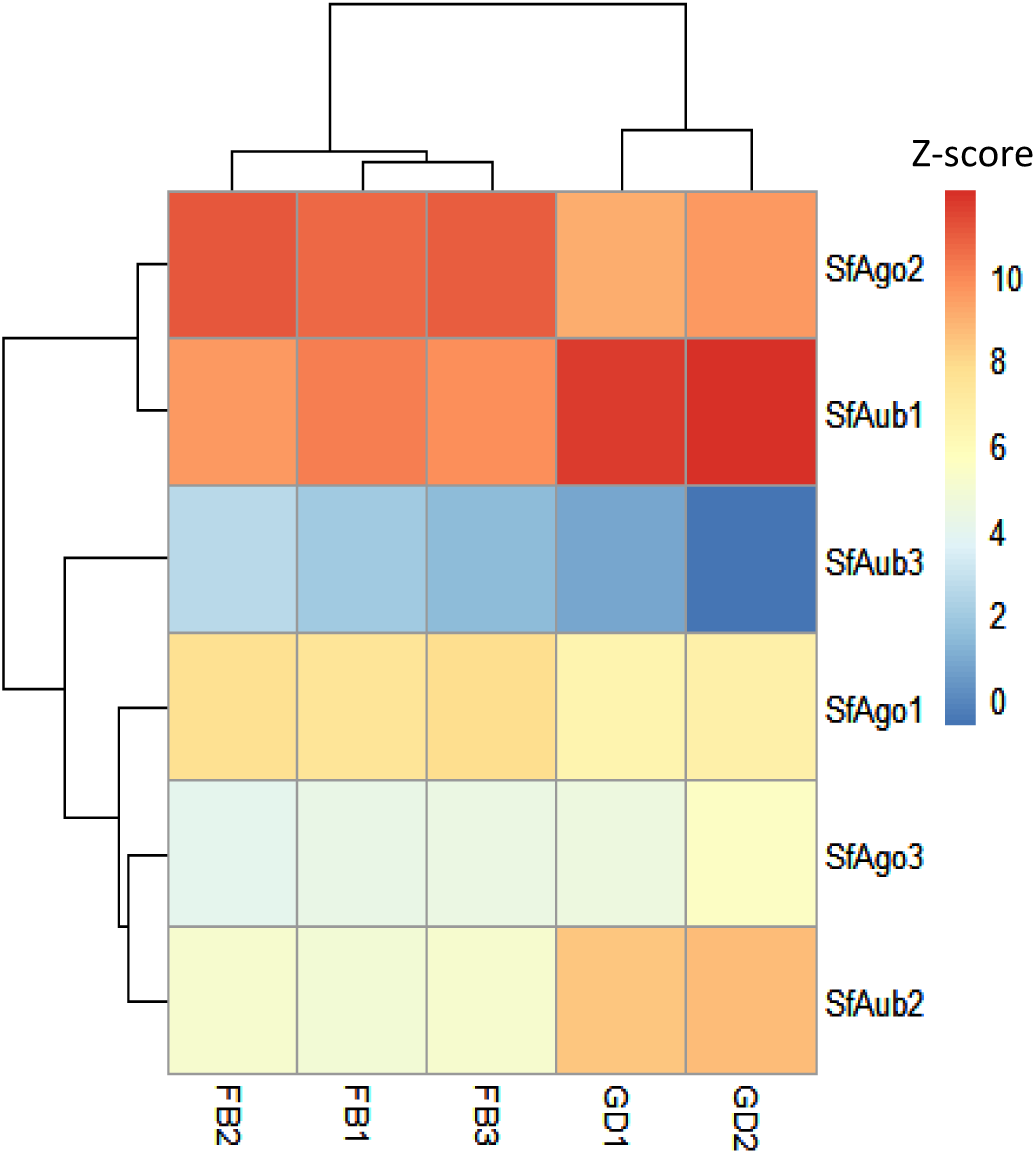
Relative levels of expression in gonadal and somatic tissues of Sf Argonaute genes. The heatmap shows the relative levels of expression of *Ago* and *Piwi* genes in the testis (GD1 & GD2 for gonads) and somatic tissue, the fat body (FB1, FB2, FB3) for each gene, with Z-score normalization.

### 3.3 piRNAs profiling in Spodoptera frugiperda

To annotate piRNAs clusters in SfC and SfR genomes, we re-used a dataset of sRNA Seq published in the frame of a previous paper describing microRNAs ^28^. Briefly, sRNAs were extracted from 4^th^ instar larvae of the lab corn strain fed on corn (CC) or rice plants (CR) as well as larvae of the lab rice strain fed on corn (RC) and rice (RR) - in duplicate. Additionally for the corn strain, sncRNA were extracted from three developmental stages, egg, larvae, pupae fed on artificial diet (CAD), Poitout (this study). The 11 libraries were analyzed by Illumina sequencing as described in ^28^. Size profiling of sRNA sequences mapping to the reference genomes (either from SfC or from SfR) and comprised between 18nt to 39nt was performed according to material and methods, the results are shown on Supplementary Figure 3. The piRNAs sequences have a size range between 24 and 39 nucleotides (nt) and include a major peak between 26-29nt and a minor peak around 33 nt. To identify piRNAs showing a ping-pong signature, we searched for piRNAs mapping to the genome on opposite strands and overlapping on 10 nucleotides using the script Ping-pong pro. We found 12935 loci to which piRNAs with ping-pong signatures mapped, among which 11185 had a score comprised between 0.5 and 1, Supplementary Table 3.

### 3.3 piRNA clusters in SfC and SfR

The pool of SfC piRNAs and of SfR piRNAs of size 24 to 39 nt corresponding to 109,416,742 and 74,370,403 sequences were mapped to the two reference genomes v5.0 and v3.0 for the corn and the rice strain, respectively using the software bowtie1. Sixty-three % of the reads from SfC gave rise to one or more alignment on the reference genome using the software Bowtie. The remaining 36.71% of reads which failed to align may map to the W chromosome which is absent in our genome assemblies built from males (ZZ). piRNAs clusters were detected using the software Shortstack with default parameters (See Material and Methods). As shown on Table 1, 521 and 628 piRNA clusters with more than 1000 reads aligned were detected in SfC and SfR genomes, respectively. The complete list of clusters with their genomic position and the sequence of the the most abundant small RNA expressed are available on Supplementary Table 4. 988 clusters had a size range of 1kb-16kb in the corn strain, 360 clusters had a size range of 1kb-7kb in the rice strain.

**Table 1.** Distribution of piRNA clusters in the genomes of SfC and SfR.

| Strain of *S. frugiperda* | SfC | SfR |
| --- | --- | --- |
| Number of clusters with more than 1000 reads aligned | 521 | 628 |
| Number of clusters larger than 1kb | 988 | 360 |
| Clusters on + strand, % | 45.8 | 46.2 |
| Clusters on – strand, % | 42.7 | 50.1 |
| Dual clusters, % | 11.5 | 3.7 |
| Total number of clusters detected | 8,889 | 3,456 |

### 3.3 Long non-coding RNAs mapping to TE, putative piRNA precursors

Long non-coding RNAs (LNCRNAs) transcribed from piRNA clusters can be precursors of piRNAs. We identified 5755 LNCRNA from contigs assembled from a set of RNA-seq libraries ^43, 59^ from various tissues of the insect - gonads, hemocytes and fat body- according to Materials and Methods. Among them, 320 show 98% identity over 80% of their length to TE sequences of various orders and can correspond to transcripts of TEs possibly embedded into piRNA clusters (Supplementary Figure 4).

### 3.3 Targets of piRNAs

piRNAs are well known regulators of TE expression, they can also act on gene expression in the germline and in the soma. We first annotated and compared TE content in the two *Sf* strains. Then, we mapped *Sf* piRNAs dataset on TE consensus or gene transcripts, to identify their targets.

#### 3.3.1 Annotation of transposable elements in SfC and SfR genomes

Using the TE denovo package of the REPET pipeline ^53,54^, we identified and classified repeated sequences in the reference genome of of the corn strain SfC v5.0 ^44^ and of the rice strain SfR v3.0 ^60^ whose assembly quality is equivalent and less fragmented than in our first reference genomes ^38^ to obtain comparable quality of TE annotation (supplementary Table 5 for statistics of genome assembly). We obtained 2171 repeats consensus in the corn strain and 2032 in the rice strain, with more retrotransposons (959 families in SfC, 872 in SfR) than DNA Tns (858 families in SfC, 821 in SfR) in both strains, as shown on Table 2.

**Table 2.** TE annotation of the two strains genomes sfC and SfR with the REPET pipeline.

| Genome assembly : | *SfCv5*<br>PACBIO Illumina<br>1000 scaffolds<br>N50=0.9Mb | *SfRv3*<br>PACBIO Illumina<br>1054 scaffolds<br>N50= 1.13Mb |
| --- | --- | --- |
| Pipeline | REPET v2.5 |  |
|  | TE DENOVO |  |
| RNA TE | 959 (44.17%) | 872 (42.91%) |
| DNA TE | 858 (39.52%) | 821 (40.40%) |
| Other repeats | 354<br>(16.31%) | 339<br>(16.69%) |
| Total consensus | 2171 (100.00%) | 2032<br>(100.00%) |
| Pipeline | TE ANNOT |  |
| Cumulative coverage | 124313817 bp<br>(113,077,456 bp) | 120296323 bp<br>(110,035,603 bp) |
| Coverage percentage | 32.34%<br>(29.42%) | 31.67%<br>(28.96%) |
| Families with full length fragments | 1892<br>(1313) | 1830<br>(1264) |

Comparison of TE diversity found in SfC and SfR shows a similar pattern (Figure 3 A and B). The complete list of repeats families and their classification according to ^55^ are available in Supplementary Table 6. A first round of TE annotation of *Sf*C and *Sf*R genomes using these 2171 and 2032 repeats consensus, respectively, led to a global repeat coverage of 29.42% and 28.96% of the two strains genomes. To optimize TE detection and identify those showing strain specificity, we performed a second round of TE annotation using a pool of the TE families found in the two strains genomes and having full length fragments in the output of the first annotation round (*i.e.* 1313 plus 1264 families). This led to coverages of 32.34% and 31.67% in *Sf*C and *Sf*R respectively (Table 2, Supplementary Table 7). The coverage of each order of TE in SfC and SfR, displayed on Figure 3C and D, shows similar patterns.

**Figure 3.**
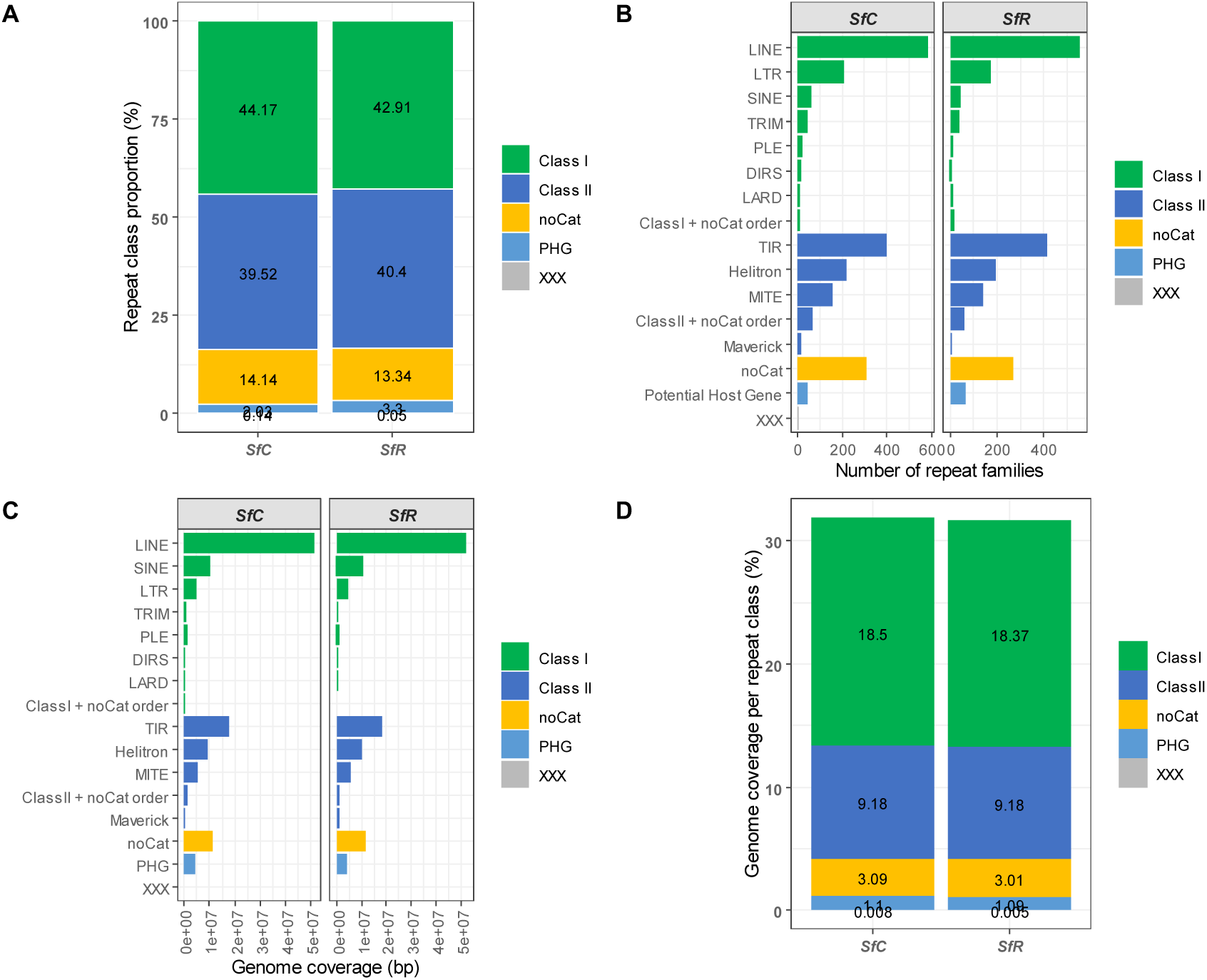
Diversity and coverage of transposable elements in the two strains genomes. A. Proportion of TE class in SfC and SfR. B. The histograms show the number of tranposable elements families in SfC (on the left, n=2171) and in SfR (on the right, n=2032) belonging to different orders according to ^55^. PHG: Potential Host Gene. noCat (repeated sequence not classified at class and order levels). XXX (sequence not classified at class level and with potential several orders) C. The histograms show the percentage of genome covered by different TE orders according to ^55^. D. proportion of the genome covered by the different repeats classes.

#### 3.3.1 piRNAs regulating TEs

To identify the TE regulated by piRNAs, we searched for piRNAs showing homology to TEs. For that purpose, sequences of size range 24nt to 39nt from SfC (109,416,742 sequences) were pooled as well as those from SfR (74,370,403 sequences) and were aligned using bowtie 1 on TE consensus sequences identified in SfC or SfR, respectively. The percentages of reads uniquely mapped on DNA TE or RT TE were calculated and shown on Figure 4A, the same analysis was performed with piRNA sequences mapping to several locations on consensus sequences (Figure 4B). The detailed counts of piRNAs mapping to the different TE consensus can be found on Supplementary Table 8. In SfC, one of the RT consensus was the most abundantly targeted transposon family, whereas in SfR reads were more evenly distributed between DNA and RT TEs. This pattern was observed both with unique and multiple mappers piRNAs sequences. The differences between histograms highlight variation in the regulation of TEs between the two strains, either because some TE can be specific of one strain or because some may be more active in one strain compared to the other.

**Figure 4.**
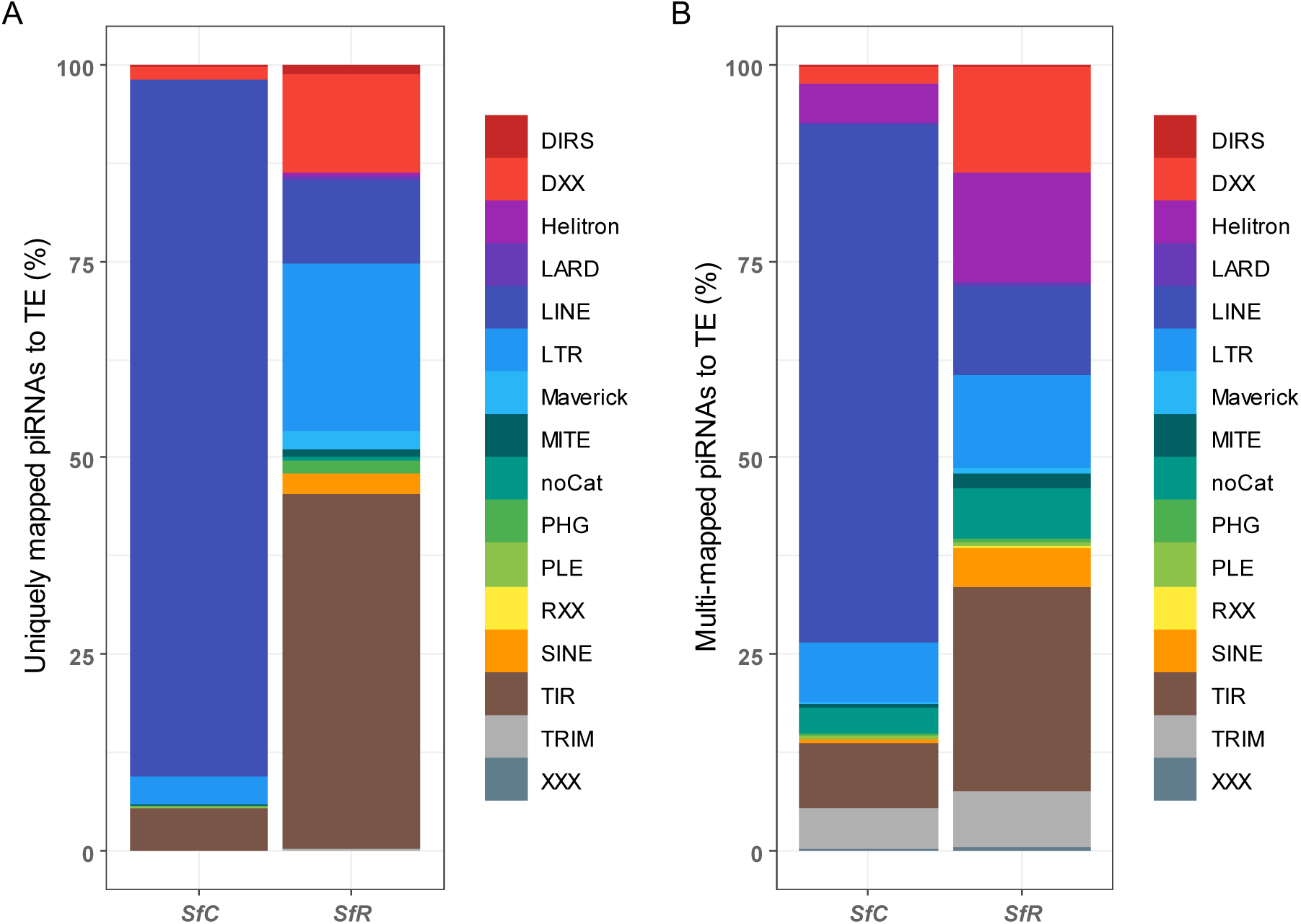
Transposable elements targets of piRNAs. piRNA were mapped to TE consensus and reads mapping uniquely (A) or at multiple location (B) were counted. The histograms show the percentage of reads mapping to different TE orders in SfC and SfR.

#### 3.3.1 piRNAs regulating cellular genes

##### 3.3.1.1 Mapping on OGS6.1

PiRNAs were mapped to the coding sequences of *S. frugiperda* genes (OGS6.1). 14577 genes out of 21830 were targeted by one or more piRNAs. 2589 genes out of 21830 were targeted by 100 piRNAs sequences or more (14577 by 1 piRNA or more). We tested whether, among them, the 2059 genes showing gene ontology (GO) annotation were enriched in some specific GO terms compared to the reference set containing 9628 genes. Ninety-five terms showed specific and statistically significant enrichment, among which the top hits were terms related to the biological process translational elongation, formation of cytoplasmic translation initiation complex, fatty acid beta-oxidation and the molecular functions structural constituent of ribosome, ATP binding, proton-transporting ATPase activity, fatty acid binding plus transposase activity (Supplementary table 9, Supplementary Figure 5).

##### 3.3.1.2 Attempt to identify a piRNA regulating sex-determination in Spodoptera frugiperda

In *Bombyx mori*, a piRNA is known to regulate sex determination by preventing the protein MASC to induce the splicing pattern of double-sex mRNA specific to the male developmental cascade ^35^. It acts by inducing a cleavage of *Masc* mRNA. Regulation of sex determination by a piRNA is not the only mechanism triggering sex determination in Lepidoptera ^61^, however involvement of the gene *Masc* seems a general feature in this insect order. In the case of *S. frugiperda*, annotation of piRNA clusters (see previous section) revealed a cluster encompassing the second exon of a gene showing homology to *masculinizer* 1 from *Helicoverpa armigera* (Figure 5 A). This piRNA cluster shows a ping-pong signature (Figure 5 B). From the sRNA-Seq data obtained from developmental stages, the candidate for pi-masc is expressed to 28 copies in eggs, 2 in larvae, 0 in pupae whereas the candidate for pi-fem is expressed to 588 copies in eggs, 10 in larvae and 0 in pupae. By mapping of RNA-Seq from testis, we observed two isoforms for *Masc*, one with all 10 exons, one with the two last exons.

**Figure 5.**
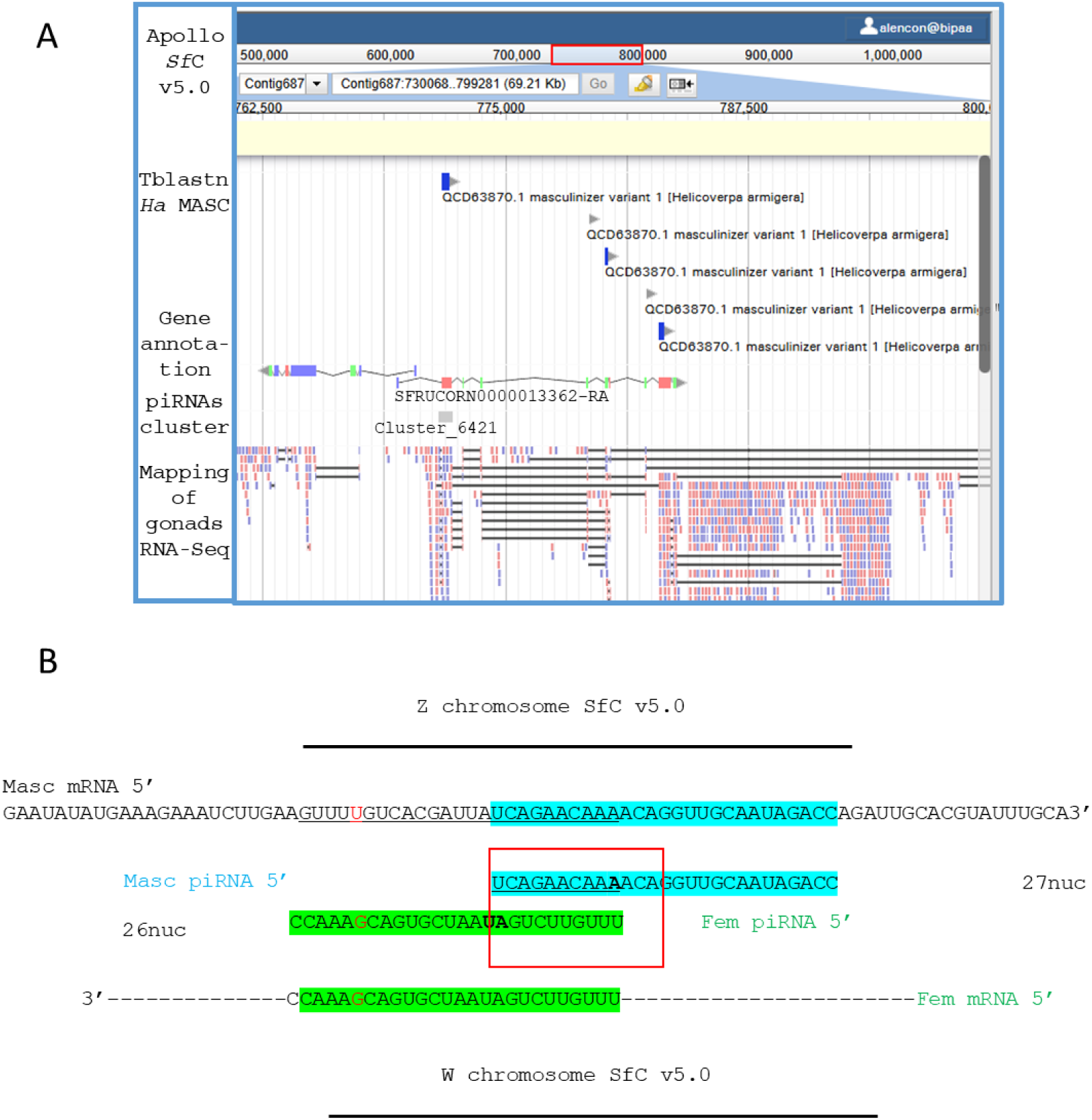
Cluster of piRNAs on *masc* gene A. *S. frugiperda* gene models matching to Masculinizer proteins from *Helicoverpa armigera* or *Sf*9 cells were identified. A piRNA cluster maps to the second exon of Sf*Masc*. B. A pair of piRNAs expressed by the cluster shows a ping pong signature.

## 4. Discussion

In this paper, we described the genes and the genomic loci involved in biosynthesis of piRNAs in *S. frugiperda* as well as the TE or genes regulated by these piRNAs. This led to two main findings i) the discovery of two new *Piwi* genes compared to the Lepidopteran model *Bombyx mori*, these two genes being conserved in the genus *Spodoptera* ii) the fact that the genomes of the two host-plant strains of *S. frugiperda*, despite of a very similar content of transposons diversity and coverage, show different patterns of piRNAs-regulated TE, suggesting a difference in transposons activity.

The phylogenetic analysis of insect Argonaute proteins showed a 1:1 orthology relationship for *Ago*1, *Ago*2 and *Ago*3 between *Spodoptera* species and *Bombyx mori*, but the existence of three *Aub* genes in the genus *Spodoptera* compared to one in *Bombyx mori* and two in *Drosophila*. In their study focused on the role of PIWI proteins in response to baculovirus infection ^30^, Xia *et al.* in 2023 found five paralogs of *Aub* in the genome assembly of an invasive population of *Spodoptera frugiperda* ^62^. By phylogenetic analysis, we show here that two of these paralogs cluster with our AUB2 and AUB3 and that they probably result from artefactual duplication during the genome assembly process due to high heterozygosity of invasive population, since the genome size 486Mb is larger than expected ^62^. Our genome assemblies were made from lab colonies of *Sf*C and *Sf*R strains originating from native area, they have a size close to expected by flow cytometry (for SfC, 1C = 394.4 +/- 0.2 Mb (N=2), Spencer Johnston, personal communication). We conclude that *Spodoptera frugiperda* has three *Aub* genes rather than five. Gene copy-number variation can occur by tandem duplications, retro-position or unequal crossing over. In our chromosomal genome assembly v7.0 ^63^, *Aub*1 is located on chromosome 11 like *Ago*1 while *Aub*2 and *Aub*3 are located on chromosome 10, like *Ago*2, at a distance between 0.5 and 2Mb from each other, it is thus unlikely that they have arisen by tandem duplication.

In *Bombyx mori*, the two *Piwi* genes, *Ago*3 and *Siwi* are highly expressed in gonads and their mRNAs can be detected in other somatic tissues in larvae or pupae ^64 65 66 67^. By RNA-seq analysis, we showed that on the contrary *Sf Ago*3 is expressed at similar level in gonads during spermatogenesis and in a somatic tissue, the fat body. However, like in *Bombyx, Sf Aub*1 (ortholog of *Bombyx Siwi*) and *Aub*2 mRNAs are more abundant in gonads compared to fat body where they can be readily detected. *Aub*3 is expressed at very low level in these two tissues.

This overexpression of the new gene *Aub*2 in gonads is in accordance with a function in the control of TEs in the germline - the primary role of piRNA pathway. In *Drosophila*, there are two *Aub* genes, one ortholog to *Siwi* (*SfAub*1) encodes AUB, a cytoplasmic protein involved in post-transcriptional silencing of TE, the other encodes PIWI, a nuclear protein responsible for transcriptional silencing of TE. *Bombyx mori* lacks functional homolog of PIWI ^64^ and thus could lack transcriptional regulation of TE. However ^68^ interactions between AGO3/SIWI and the *Bombyx* proteins HP1a and HP1b have been shown suggesting that *Bombyx* PIWI proteins could play the role of transcriptional regulators. *Sf* AUB2 could be a functional homolog of *Drosophila* PIWI. Alternatively, AUB2 could replace AGO3 and compensate for its low expression in gonads. We are currently characterizing a KO mutant of this gene to assess its function (Gasser *et al.*, in prep.).

The very low expression of *Aub*3 in gonads suggests that it evolved a new function. As reviewed in ^69^, diversification of *Piwi* genes in insects led to diversification of the functions of PIWI proteins. For instance, in the pea aphid ^18^, the eight *Piwi* genes have evolved specialization of their expression pattern, two of them, *Piwi*2 and 6 are expressed in the germline, two others, *Piwi*5 and 3 in the soma, *Piwi*5 is more expressed in males, *Piwi*6 and *Ago*3 are expressed in sexuparae. In mosquitoes, the main role of PIWI proteins and piRNAs is the control of viral infections in different somatic tissues and cell lines (^69^ for review). In Lepidopteran cell cultures from *B. mori* and *Trichoplusia. ni* Santos *et al.*, showed by RNA interference that PIWI proteins are involved in antiviral immunity ^70^. In the study that we mentioned in the beginning of the discussion section^30^, Xia *et al*. showed using dsRNA silencing in *Sf*9 cells that their Piwi-like-4-5 protein (Our AUB3) significantly promotes the Baculovirus Ac MNPV replication and thus acts as a proviral factor like SIWI and AGO3 in *Bombyx mori*. On the contrary their Piwi-like-1 (Our AUB2) and their Piwi-like-2-3 (Our AUB1) have an antiviral activity against Ac MNPV. In their study these functions related to viral infection seem to be independent on viral piRNAs, they involve only PIWI proteins. This study does not provide any data on the *Piwi* gene expression level in *Sf*9 cells.

Biogenesis of piRNAs needs transcription of genomic regions enriched in remnants of transposable elements by RNA-PolII in Drosophila recruited by the HP1 homolog Rhino ^71^. The processing of these transcripts to mature piRNAs involves various enzymes PIWI, Zuchini and a 3’ to 5’ trimming activity (details reviewed in ^72^). To annotate these piRNA clusters in the two *S. frugiperda* genomes, we relied on sRNA-Seq data from 11 libraries ^28, this study^. The piRNA size profiling highlighted in all libraries, a major peak in the size range of 26-29 nt and a secondary peak of 32-33 nt (Supplementary Figure 3). This secondary peak can correspond to an intermediate of biosynthesis or to mature piRNAs produced by the sur-numerary PIWI proteins found in *S. frugiperda* like in *Drosophila* where each PIWI family member was shown to bind different populations of small RNAs ^21^. Since we do not have antibodies against *Sf* PIWI proteins, we could not purify piRNAs further. However from the pie-chart representing the proportion of small RNAs mapping to various genomic references (Supplementary Figure 4), there was a low proportion of sRNAs mapping to rDNA whereas the vast majority of cellular RNA consists of ribosomal RNA (rRNA) - ∼ 80 to 90% of total RNA for most cells^73^, showing that contamination of our samples with sRNA sequences resulting from degradation of large RNAs was minimal. We found 500-600 clusters with more than 1000 piRNA reads aligned and between 400-1000 clusters larger than 1 kb scattered on most of the genomic scaffolds in *Sf*C and *Sf*R, respectively. In *Bombyx mori* genome, depending on the methods and sRNA-Seq libraries used to annotate them, between 560 and 965 clusters were described ^74, 75^, which corresponds to a similar magnitude order. This is more than in *Drosophila* where 142 genomic locations were identified as sites of abundant piRNA generation ^21^. We found both unidirectional and bi-directional piRNA clusters in *S. frugiperda*. In *Drosophila* piRNA clusters transcribed on both DNA strands are expressed in the germline^21^. Because TEs in piRNA clusters can accumulate mutations that negate their transposition activity, we searched for long non-coding RNAs homolog to TE from a dataset of RNA-Seq ^43,59^. We identified 325 such transcripts as putative piRNA precursors. The epigenetic marks associated to piRNA clusters vary : In *Drosophila*, piRNA clusters are embedded in heterochromatin regions^71^. In *Bombyx*, they are mostly found in euchromatin except telomeric ones which are heterochromatic^75^. Accordingly, *Drosophila* Piwi interacts with HP1^76^, whereas *Bombyx* Siwi interacts both with a Histone H3K4 methyl transferase^77^ and with HP1^68^.

We found 32.34% and 31.67% of the genomes of the two strains SfC and SfR corresponding to repeated sequences, slightly more than in the first genome paper ^27^ (29.16% - 29.10%, respectively) reflecting the better contiguity of our genome assemblies. The two strains share a similar pattern of TE diversity, with a predominance of Non-LTR retrotransposons and SINES, like in *B. mori* ^78^. However, we found variation between the two strains when we compared the patterns of TE targeted by piRNAs showing that active transposons differ between them. Initially described as host-plant strains due to their association to different plant ranges ^79^, the evolutionary status of the two strains has recently been clarified by population genomics analysis : the whole genome sequences of larvae collected on corn or rice cluster in two genetically distinct groups ^26^ with very few hybrids. They are in the process of incipient speciation and display traits acting as reproductive barriers, reviewed in ^80^. In *Drosophila melanogaster*, there are strains called “P strains” with the P transposable element and some which do not contain it called “M strains” ^81^. in P strains, P element-derived piRNAs are transmitted from the mother through the oocyte cytoplasm. The transmitted piRNAs then silence the P element. If no piRNA is transmitted from the mother, the P element is not repressed. Consequently, a P male crossed with an M female will have a dysgenic offspring, with abnormally small gonads, increased mutation rates, frequent sterility. This phenomenon, resulting from the fact that the offspring have the P element but no silencing through maternal piRNA, is known as “hybrid dysgenesis” ^82^. Similarly in *S. frugiperda* TE families active in one strain and not in the other could impair the fitness of hybrids and reinforce reproductive isolation. We will test this hypothesis in future.

In addition to TE, we identified 2589 genes which were targeted by at least 100 piRNA sequences, likely regulated by piRNAs and PIWI proteins. A GO term enrichment analysis of them showed that among the most enriched biological processes figure translation elongation, macromolecule catabolic process, protein phosphorylation, regulation of mRNA stability, transposition. The fact that among regulated genes, some are related to transposition was expected since our gene annotation was performed without hiding repeated sequences and contains some TE genes. Among the most enriched cellular components terms, the eukaryotic translation initiation factor 3 complex was found as well as eukaryotic 43S and 48S preinitiation complexes. PIWI proteins are known to be involved in translation initiation in germ cells in *Drosophila* ^83^ as well as in mouse during spermatogenesis ^84^, suggesting that this role of PIWI-piRNAs in translation initiation is conserved in *S. frugiperda*. One of the first genes shown to be regulated by a piRNA is the sex determining gene *Masc* in *Bombyx mori* ^35^. Since that time, alternative mechanisms to trigger the sex determining cascade have been discovered in Lepidoptera (reviewed in ^61^). In the frame of the annotation of piRNA clusters in *S. frugiperda*, we discovered a piRNA cluster in the second exon of *Masc*, coding for a pair of piRNAs showing the ping-pong signature. We are limited at present to study more deeply these piRNAs as candidates for sex-determining gene regulation because the W is absent in our genome assemblies and we could not identify a *fem* precursor transcript in the transcriptomic resources available today. However, because sex determining piRNAs are species specific and small size RNA sequences, they could be of interest in the frame of development of new biological control means for this insect pest populations.

## Acknowledments

We thank Johann Confais at URGI INRAE Versailles for advises with the REPET pipeline. This work was supported by two grants from the Plant Health and Environment Division of INRAE: a “HOLOCEN” grant awarded to Emmanuelle d’Alençon and a “CENTROVIR” grant awarded to Isabelle Darboux. Mélanie Gasser was awarded a post-doctoral fellowship through the ANR-DFG project ORIGINS, ANR-20-CE92-0018-01.

## Data availability

SfC v5.0 genome assembly is available at ENA as PRJEB96863 (ERP179489). Its is also available as well as v3.0, the genome assembly of SfR, with gff files corresponding to TE and piRNA clusters annotation on the bioinformatic platform BIPAA at, respectively : https://bipaa.genouest.org/sp/spodoptera_frugiperda_corn/ https://bipaa.genouest.org/sp/spodoptera_frugiperda_rice/.

