## Supplementary figures for "A characterization of piARNs, their biogenesis and their targets in *Spodoptera frugiperda* (Lepidoptera, Noctuidae)"

Supplementary Figure 1. Phylogeny and corresponding IDs of the piwi genes identified in native populations ofSfC (this study) and those identified in invasive populations in China by Xia et al., 2023

Supplementary Figure 2. PCA and MA plot obtained during the RNA-Seq analysis published in our previous paper (Gimenez et al., 2025) and re-used in this study

Supplementary Figure 3. Size profiling and pie-chart representing the proportion of sRNA sequences mapping to different genomic references.

Supplementary Figure 4. LNCRNAs mapping to transposable elements and DNA repeats in SfC

Supplementary Figure 5. GO term enrichment analysis of the coding genes regulated by piRNAs

### Sup. Fig. 1

Phylogeny and corresponding IDs of the piwi genes identified in native populations of SfC (this study) and those identified in invasive populations in China by Xia et al., 2023

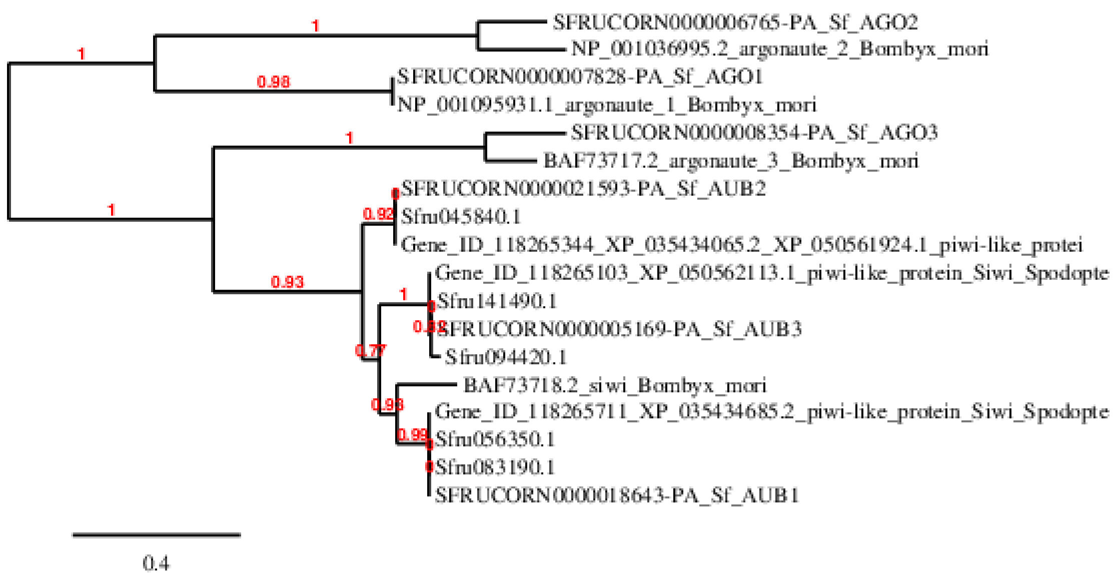


| Gene name | ID OGS6.0<br>SfCv5.0 | ID OGS6.1<br>Transferred in SfCv6.0 | Gene_ID in [1] | ID in [2] |
|---|---|---|---|---|
| SfAgo1 | SFRUCORN0000007828-RA | SFRUCORN610000007828-RA | ND | ND |
| SfAgo2 | SFRUCORN0000006765-RA | SFRUCORN610000006765-RA | | |
| SfAgo3 | SFRUCORN0000008354-RA | SFRUCORN610000008354-RA | | |
| SfAub1 | SFRUCORN0000018643-RA | SFRUCORN610000018643-RA | 118266049* Siwi2<br>118265711 siwi3 | Sfru056350.1<br>Sfru083190.1 |
| SfAub2 | SFRUCORN0000021593-RA | SFRUCORN610000021593-RA | 118265344 Siwi1 | Sfru045840.1 |
| SfAub3 | SFRUCORN0000005169-RA | SFRUCORN610000005169-RA | 118265181* Siwi4<br>118265103 Siwi5 | Sfru094420.1<br>Sfru141490.1 |

- * suppressed in Genbank
- In bold, Two transcripts XP_050561924.1 and XP_035434065.2

1. Xia, J.M., Fei, S.G., Wu, H.Y., Yang, Y.F., Yu, W.S., Zhang, M.M., Guo, Y.Y., Swevers, L., Sun, J.C., and Feng, M. (2023). The piRNA pathway is required for nucleopolyhedrovirus replication in Lepidoptera. Insect Science *30*, 1378–1392. 10.1111/1744-7917.13160.

2. Xiao, H., Ye, X., Xu, H., Mei, Y., Yang, Y., Chen, X., Yang, Y., Liu, T., Yu, Y., Yang, W., et al. (2020). The genetic adaptations of fall armyworm Spodoptera frugiperda facilitated its rapid global dispersal and invasion. Mol Ecol Resour *20*, 1050–1068. 10.1111/1755-0998.13182.

PIWI protein sequences described in Xia et al., 2023 and identified in Xiao et al. 2020

>Gene_ID_118265344_XP_035434065.2_XP_050561924.1 piwi-like protein Siwi [Spodopte
MADSGKKKTKRAGKKFKKNAPPPLKLPTEQPGPSGSDPKSPPSAGSSVQSPPSAGPSVQSPPSAGPSVQS
PPSAGPSVQSPTAAGPGPKSSSTAGPALKRTPAKETGLQSTPAGGHGSKTPGPQQKSRPAEKLSLQSPSA
PGPSTQEPPVWPPPSQKKLSLPQESKAEHRAAPSVREQPPGDVQIEQKLQSIHIGEKPTESGSKSKGASI
LRTRPIELDSKKGMSGTLVDVKANYFTVETTPKWCLYQYHVDFSPDEDSTAVRKALMRIHAKTFGGYLFD
GTVLYTVKRLRPDPLELFSIRSNDEQRMRILIKLTCDVAPGDYHYIQIFNIIIRKCFQILKLQLMGRDYF
DPLAKIDIPEYRLQIWPGYKTTINQYEDRLLMVTEITHKVLRLDTVLQMICDYADSKGANYRKIFLEDIV
GKIVMTIYNKKTYRVDDVAWQVTPKDTFKIRDGSITYVDYYKNKYNLIIRDLQQPLLISRSKPRDIRAGM
PELVYLVPELCRQTGLTDEMRNNFKLMKALDNYTKVGPDVRIQKLLHFNKRLTQNREVLKEMAEWSLTLS
KDLVNFKARQLPPENIYQGNNVSYQAGDTADGWTRSMRSHPLLAIAQMPSWVVISPDRLQRDTEGFIELI
TKTAYSVGLRMPKPELVLIKYDSAMEYASQCETAIARKNPAMILCVLTRKVSDRYEAIKKKCTVDRAVPT
QVVCARNMTATSAMSIATKIAIQINCKLGGAPWRVTIPLKGCLVIGYDVCHDTRSKEKSFGGFVASLDDH
LTKFYSAVNSHSSGEELSNHMGFNMETTLTKYKMKNGRLPDRIFIYRDGVGDGQIPYVYSHEVVQIKKKL
AELYGGAPVKMAFIIVSKRINTRIFLNRGQGENPRPGTVIDDVITLPERYDFYLVSQNVRNGTISPTSYN
IIENTTGMHPDRLQWLTYKLTHMYFNCSAQVRVPAVCQYAHKLAFLAANSLHSQPHYTMTDTLYFL

>Gene_ID_118265711_XP_035434685.2 piwi-like protein Siwi [Spodoptera frugiperda]
MSQPGGRGWGRGRAGRGGDNAGPPPRRPGEPARPVPTQQPVPTGPRPQPPTPWGPPTAQAGRASHRSTPS
THQEHPGDVDVQQRMQAMQIAGTSQPTAAGEPGSAVIGRGSRRGGGRVLPEQMTILRTRPSTIDSKKGTS
GTVIDLRSNYFMVKTTPQWRLYQYHVDFSPDEDSTFIRRALMRVHAKTLGGYLFDGSVLYTVKRLHPDPL
ELYSDRKTDDQRMRILIKLTCDVAPGDYHYIQIFNIIIRKCFQILDLQLMGRDYFDSEAKVDIPEYRLQI
WPGYKTTINQYEDRLLMVTEITHKVLRLDTVLQMLSDYAASKGGNYKKIFLEDIVGKIVMTTYNKKTYRV
DDVSWQVSPKSTFKMRDETITYVDYYRKKYKINIQDTAQPLLISRSKPRDIRAGMPELVYLVPELCRQTG
LTDEMRSNFKLMKALDAHTKIGPDVRIQKLMAFNQRLTRNPEVVKEMGEWDLTLSNELIKIKGRQLPPEN
IVQGNNNRYPAGHTTDGWTQDMRSKPLLEIAQLSSWVVITPERQRRETEGFVDLIMKTAGGVGFRMPRPE
VVTIRNDSSMEYANACENSIARKNPAMLLVVLGRKVTDRYEAIKKKCTVDRAIPTQVVCARNMTSKSAMS
IATKVAIQINCKLGGAPWRVDIPLQNILVIGYDVCHDTRSKEKSFGGFVASLDRYLTRYFSAVNSHTSGE
ELSSHMSFNVGSALKKFREVNGTLPSRVFIYRDGVGDGQIPYVHSHEVGEIKKKLAEIYAGEEYKMAFII
VSKRINTRIFLDRGRGENPRPGTVVDDVVTQPERYDFYLVSQNVREGTISPTSYNVIEDSTGMHPDRMQW
LTYKMTHLYYNCSSQVRVPAVCQYAHKLAFLAANSLHSQPHYSLTDTLYFL

>Gene_ID_118265103_XP_050562113.1 piwi-like protein Siwi [Spodoptera frugiperda]
MPKKRLKNRAGKTSKNASQSSRKSGDPPTCPQSPQTPTSPSAWSPRTSQSRKSTTSKAQTVSSSPTSPSI
PSSQTTLTSTLPHGTPTSQEGRRSSKQSRLQEQSEDVPKQLQRMSIDKPTSRSNISSKPGPSSSGRGHGA
RPNTEQRPALRTKPSTISSKKGSTGNAVNLIANYFAVETTPHFRMYQYHVDFSPDEENTFVRKSLMRIHQ
KVLGWHVFDGSVIYSMKRLHPDPTEFYSDRKSDKQKMVITVKLIGDVAPGDYHYIQIFNIILRKCFLAMD
LQLMGRDYFDPTAKIDIPEFRLQIWPGYKTTINQYEDKLLMVAVIIHKVLRLDTVLQMLKTYTNTKGGNY
KKCFLEDIVGKIVMTNYNRRTYRVDDVSWLTTPKSTFNLRNVKISYIEYYKKKYNINIRDVDQPLLISRS
KPRDIRAGMPELIYLVPELCRQTGLTEEMRNNFKLTKALDAHTKIGPDIRVNKLLQFNRNLTQNPEVIKE
LNTCHLTFSKQLMRIKGRQLPTENIVQGNECRYPAGDTPEGWTREVRSKPLLRISQLPSWVVITPDRARR
DTEAFLDMVIKAARDVRFMLPKPEIVTMQYDGPIDYARMLDAVIARRNPAMILCVVTRKLSDRYEAIKKK
CLVDRAIPTQVVTLRNMTSNSAMSVATKVALQMNCKLGGAPWRVEIPLQNILVVGYDVCHDTRSKQKSFG
GFVASLDRHMTRYYSAVNSHTSGEELSNHMGFNTASALKKYREVNGVLPHRIFIYRDGVGDGQIPYVHTH
EVAEINKALRQLYGPDEFKMAFTIVSKRINTRFFLEQGSRCVNPRPGTVVDDVVTLPERYDFFLISQNVR
DGTITPTSYNVISDSTGLHPDRMQWLTYKLTHMYFNCSAQVRVPAVCQYAHKLAFLAANSLHSQPHNSLT
DTLYFL

### Proteins in Xia et al., 2023 which have been suppressed

Gene_ID_118266049_XP_035435290.1_piwi-like protein Siwi
Same as Sfru083190.1

MSQPGGRGWGRGRAGRGGDNAGPPPRRPGEPARPVPTQQPVPTGPRPQPPTPWGPPTAQA
GRASHRSTPSTHQEHPGDVDVQQRMQAMQIAGTSQPTAAGEPGSAVIGRGSRRGGGRVLP
EQMTILRTRPSTIDSKKGTSGTVIDLRSNYFMVKTTPQWRLYQYHVDFSPDEDSTFIRRA
LMRVHAKTLGGYLFDGSVLYTVKRLHPDPLELYSDRKTDDQRMRILIKLTCDVAPGDYHY
IQIFNIIIRKCFQILDLQLMGRDYFDSEAKVDIPEYRLQIWPGYKTTINQYEDRLLMVTE
ITHKVLRLDTVLQMLSDYAASKGGNYKKIFLEDIVGKIVMTTYNKKTYRVDDISWQVSPK
STFKMRDETITYVDYYKKKYKINIQDTAQPLLISRSKPRDIRAGMPELVYLVPELCRQTG
LTDEMRSNFKLMKALDAHTKIGPDVRIQKLMAFNQRLTRNPEVVKEMGEWDLTLSNELIK
IKGRQLPPENIVQGNNNRYPAGHTTDGWTQDMRSKPLLEIAQLSSWVVITPERQRRETEG
FVDLIMKTAGGVGFRMPRPEVVTIRNDSSMEYANACENSIARKNPAMLLVVLGRKVTDRY
EAIKKKCTVDRAIPTQVVCARNMTSKSAMSIATKVAIQINCKLGGAPWRVDIPLQNILVI
GYDVCHDTRSKEKSFGGFVASLDRYLTRYFSAVNSHTSGEELSSHMSFNVGSALKKFREV
NGTLPSRVFIYRDGVGDGQIPYVHSHEVGEIKKKLAEIYAGEEYKMAFIIVSKRINTRIF
LDRGRGENPRPGTVVDDVVTQPERYDFYLVSQNVREGTISPTSYNVIEDSTGMHPDRMQW
LTYKMTHLYYNCSSQVRVPAVCQYAHKLAFLAANSLHSQPHYSLTDTLYF

Gene_ID_118265181_XM_035577959.1 piwi-like protein Siwi
Same as Sfru094420.1

MPKKRLKNRAGKTSENASQSTRRSGDPPTCPQSPQTPTSPQSPQ
TPTSPSAWGPHTSQSRKSTTSKAQPVSSSQQSPTSPSPPKSPIYPTSPLPRGTPTSQE
GRRSSKQSEDVPRQLQKVKSIDKPTSRSNISSKPGPSSSGRGHGGRTDTEQRPALRTK
PSTISSKQGSTGNAVNLIANYFAVETTPHFRMYQYHVDFSPDEENTFVRKSLMRIHQK
VLGWHVFDGSVLYSMKRLHPDPTEFYSERKSDKQKMILTVKLIGDVAPGDYHYIQIFN
IILRKCFLVMDLQLMGRDYFDPTAKIDIPEFRLQIWPGYKTTINQYEDKLLMVAVITH
KVLRLDTVLQMLKTYTTTKGGNYKKCFLEDIVGKIVMTNYNRRTYRVDDVSWLATPKS
TFNLRNEKISYIDYYKKKYSINIRDVDQPLLISRSKPRDIRAGMPELVYLVPELCRQT
GLTEEMRNNFKLMKALDAHTKIGPDIRVNKLLQFNRNLTQNPEVIKELNTCHLTFSKQ
LMRIKGRQLPTENIVQGGECRYPAGDTPEGWTREVRSKPLLRISLLPSWVVITPDRAR
RDTEAFLDMVIKAGRDVKFMLPKPEIVTMQYDGPIEYARMLDAVIARRNPAMILCVVT
RKLSDRYEAIKKKCLVDRAIPTQVVTLRNMTSNSAMSVATKVALQMNCKLGGAPWRVE
IPLQNILVVGYDVCHDTRSKQKSFGGYVASLDRHMTRYYSAVNSHTSGEELSNHMGFN
TASALKKYREVNGMLPHRIFIYRDGVGDGQIPYVHTHEVAEINKALRQLYGPDEFKMA
FTIVSKRINTRFFLEQGSRCVNPRPGTVVDDVVTLPERYDFFLISQNVRDGTITPTSY
NVISDSTGLHPDRMQWLTYKLTHMYFNCSAQVRVPAVCQYAHKLAFLAANSLHSQPHN
SLTDTLYFL

PIWI protein sequences described in Xia et al., 2023 and identified in Xiao et al. 2020

>Sfru056350.1 seq ID
MAGFQTSDTMPKFGHVTDLREQWLVRADGALAHRLQDREISSHLCENRYRNQQIREDFPIALNEQRDIEHEVHLRELTRRRQAEID
AEIAREMAEINLRRQRRQRSSDMLQQPSSSRAAPPAPLPVTSVPSEDINPEELGLSPTELEEMRRRLEQEEKDARLARELSHEDNN
QDALLADMRCAVEAQDGELARLLQEREYKKLQRAKEKARQKALLKKQQRMAAQQQGQAPQVDSEASSTPQATLCDAYSNPRDMIRE
NRLRLCKSVDSDLHYVEPFEKESIQSKASALQSLEMSNQYYTRQPEAGTSGSNLSHSHSQSIEDKKVPSNQSSVQHRQPPPPPTTS
KKPRFPDPVSIRPASHYSTEQTVPESIPPHNMAHSYSENNNLYSCTYPPDINPPHMASDSHDNNKRLSIISDISAEGLAKKKSKQA
DMNKKRRGCKIHGVTTGWQVGKFPSGLLYFIFDRSQLRWVCYEPPESCFGLCFEALQAVTANFLCPCFQSHNDTLICMSQPGGRGW
GRGRAGRGGDNAGPPPRRPGEPARPVPTQQPVPTGPRPQPPTPWGPPTAQAGRASHRSTPSTHQEHPGDVDVQQRMQAMQIAGTSQ
PTAAGEPGSAVIGRGSRRGGGRVLPEQMTILRTRPSTIDSKKGTSGTVIDLRSNYFMVKTTPQWRLYQYHVDFSPDEDSTFIRRAL
MRVHAKTLGGYLFDGSVLYTVKRLHPDPLELYSDRKTDDQRMRILIKLTCDVAPGDYHYIQIFNIIIRKCFQILDLQLMGRDYFDS
EAKVDIPEYRLQIWPGYKTTINQYEDRLLMVTEITHKVLRLDTVLQMLSDYAASKGGNYKKIFLEDIVGKIVMTTYNKKTYRVDDM
SWQVSPKSTFKMRDETITYVDYYRKKYKINIQDTAQPLLISRSKPRDIRAGMPELVYLVPELCRQTGLTDEMRSNFKLMKALDAHT
KIGPDVRIQKLMAFNQRLTRNPEVVKEMGEWDLTLSNELIKIKGRQLPPENIVQGNNNRYPAGHTTDGWTQDMRSKPLLEIAQLSS
WVVITPERQRRETEGFVDLIMKTAGGVGFRMPRPEVVTIRNDSSMEYANACENSIARKNPAMLLVVLGRKVTDRYEAIKKKCTVDR
AIPTQVVCARNMTSKSAMSIATKVAIQINCKLGGAPWRVDIPLQNILVIGYDVCHDTRSKEKSFGGFVASLDRYLTRYFSAVNSHT
SGEELSSHMSFNVGSALKKFREVNGTLPSRVFIYRDGVGDGQIPYVHSHEVGEIKKKLAEIYAGEEYKMAFIIVSKRINTRIFLDR
GRGENPRPGTVVDDVVTQPERYDFYLVSQNVREGTISPTSYNVIEDSTGMHPDRMQWLTYKMTHLYYNCSSQVRVPAVCQYAHKLA
FLAANSLHSQPHYSLTDTLYFL
>Sfru083190.1
MSQPGGRGWGRGRAGRGGDNAGPPPRRPGEPARPVPTQQPVPTGPRPQPPTPWGPPTAQAGRASHRSTPSTHQEHPGDVDVQQRMQ
AMQIAGTSQPTAAGEPGSAVIGRGSRRGGGRVLPEQMTILRTRPSTIDSKKGTSGTVIDLRSNYFMVKTTPQWRLYQYHVDFSPDE
DSTFIRRALMRVHAKTLGGYLFDGSVLYTVKRLHPDPLELYSDRKTDDQRMRILIKLTCDVAPGDYHYIQIFNIIIRKCFQILDLQ
LMGRDYFDSEAKVDIPEYRLQIWPGYKTTINQYEDRLLMVTEITHKVLRLDTVLQMLSDYAASKGGNYKKIFLEDIVGKIVMTTYN
KKTYRVDDISWQVSPKSTFKMRDETITYVDYYKKKYKINIQDTAQPLLISRSKPRDIRAGMPELVYLVPELCRQTGLTDEMRSNFK
LMKALDAHTKIGPDVRIQKLMAFNQRLTRNPEVVKEMGEWDLTLSNELIKIKGRQLPPENIVQGNNNRYPAGHTTDGWTQDMRSKP
LLEIAQLSSWVVITPERQRRETEGFVDLIMKTAGGVGFRMPRPEVVTIRNDSSMEYANACENSIARKNPAMLLVVLGRKVTDRYEA
IKKKCTVDRAIPTQVVCARNMTSKSAMSIATKVAIQINCKLGGAPWRVDIPLQNILVIGYDVCHDTRSKEKSFGGFVASLDRYLTR
YFSAVNSHTSGEELSSHMSFNVGSALKKFREVNGTLPSRVFIYRDGVGDGQIPYVHSHEVGEIKKKLAEIYAGEEYKMAFIIVSKR
INTRIFLDRGRGENPRPGTVVDDVVTQPERYDFYLVSQNVREGTISPTSYNVIEDSTGMHPDRMQWLTYKMTHLYYNCSSQVRVPA
VCQYAHKLAFLAANSLHSQPHYSLTDTLYFL

>Sfru045840.1
MADSGKKKTKRAGKKFKKNAPPPLKLPTEQPGPSGSDPKSPPSAGSSVQSPPSAGPSVQSPPSAGPSVQSPTAAGPGPKSSSIAGP
ALKRTPAKETGLQSTPAGGHGSKTPGPQQKSRPAEKLSLQSPSAPGPSTQEPPVWPPPSQKKLSLPQESKAEHRAAPSVREQPPGD
VQIEQKLQAIHIGEKPTESGSKSKGASILRTRPIELDSKKGMSGTLVDVKANYFTVETTPKWCLYQYHVDFSPDEDSTAVRKALMR
IHAKTFGGYLFDGTVLYTVKRLRPDPLELFSIRSNDEQRMRILIKLTCDVAPGDYHYIQIFNIIIRKCFQILKLQLMGRDYFDPLA
KIDIPEYRLQIWPGYKTTINQYEDRLLMVTEITHKVLRLDTVLQMICDYADSKGANYRKIFLEDIVGKIVMTIYNKKTYRVDDVAW
QVTPKDTFKIRDGSITYVDYYKNKYNLIIRDLQQPLLISRSKPRDIRAGMPELVYLVPELCRQTGLTDEMRNNFKLMKALDNYTKV
GPDVRIQKLLHFNKRLTQNREVLKEMAEWSLTLSKDLVNFKARQLPPENIYQGNNVSYQAGDTADGWTRSMRSHPLLAIAQMPSWV
VISPDRLQRDTEGFIELITKTAYSVGLRMPKPELVLIKYDSAMEYASQCETAIARKNPAMILCVLTRKVSDRYEAIKKKCTVDRAV
PTQVVCARNMTATSAMSIATKIAIQINCKLGGAPWRVTIPLKGCLVIGYDVCHDTRSKEKSFGGFVASLDDHLTKFYSAVNSHSSG
EELSNHMGFNMETTLTKYKMKNGRLPDRIFIYRDGVGDGQIPYVYSHELQWLTYKLTHMYFNCSAQVRVPAVCQYAHKLAFLAANS
LHSQPHYTLTDTLYFL

>Sfru094420.1
MPKKRLKNRAGKTSENASQSTRRSGDPPTCPQSPQTPTSPQSPQTPTSPSAWGPHTSQSRKSTTSKAQPVSSSQQSPTSPSPPKSP
IYPTSPLPRGTPTSQEGRRSSKQSEDVPRQLQKVKSIDKPTSRSNISSKPGPSSSGRGHGGRTDTEQRPALRTKPSTISSKQGSTG
NAVNLIANYFAVETTPHFRMYQYHVDFSPDEENTFVRKSLMRIHQKVLGWHVFDGSVLYSMKRLHPDPTEFYSERKSDKQKMILTV
KKYSINIRDVDQPLLISRSKPRDIRAGMPELVYLVPELCRQTGLTEEMRNNFKLMKALDAHTKIGPDIRVNKLLQFNRNLTQNPEV
IKELNTCHLTFSKQLMRIKGRQLPTENIVQGGECRYPAGDTPEGWTREVRSKPLLRISLLPSWVVITPDRARRDTEAFLDMVIKAG
RDVKFMLPKPEIVTMQYDGPIEYARMLDAVIARRNPAMILCVVTRKLSDRYEAIKKKCLVDRAIPTQVVTLRNMTSNSAMSVATKV
ALQMNCKLGGAPWRVEIPLQNILVVGYDVCHDTRSKQKSFGGYVASLDRHMTRYYSAVNSHTSGEELSNHMGFNTASALKKYREVN
GMLPHRIFIYRDGVGDGQIPYVHTHEMQWLTYKLTHMYFNCSAQVRVPAVCQYAHKLAFLAANSLHSQPHNSLTDTLYFL
>Sfru141490.1 1
MPKKRLKNRAGKTSKNASQSSRKSGDPPTCPQSPQTPTSPSAWSPRTSQSRKSTTSKAQPVSSSPTSPSPSIPSSQTTLTSTLPHGTP
TSQEGRRSSKQSRLQEQSEDVPKQLQRMSIGSTGNAVNLIANYFAVETTPHFRMYQYHVDFSPDEENTFVRKSLMRIHQKVLGWHV
FDGSVIYSMKRLHPDPTEFYSDRKSDKQKMVITVKLIGDVAPGDYHYIQIFNIILRKCFLAMDLQLMGRDYFDPTAKKYNINIRDV
DQPLLISRSKPRDIRAGMPELIYLVPELCRQTGLTEEMRNNFKLTKALDAHTKIGPDIRVNKLLQFNRNLTQNPEVIKELNTCHLT
FSKQLMRIKGRQLPTENIVQGNECRYPAGDTPEGWTREVRSKPLLRISQLPSWVVITPDRARRDTEAFLDMVIKAARDVRFMLPKP
EIVTMQYEGPIDYARMLDAVIARRNPAMILCVVTRKLSDRYEAIKKKCLVDRAIPTQVVTLRNMTSNSAMSVATKVALQMNCKLGG
APWRVEIPLQNILVVGYDVCHDTRSKQKSFGGYVASLDRHMTRYYSAVNSHTSGEELSNHMGFNTASALKKYREVNGVLPHRIFIY
RDGVGDGQIPYVQTHEVAEINKALRQLYGPDEFKMAFTIVSKRINTRFFLEQGSRCVNPRPGTVVDDVVTLPERYDFFLISQNVRD
GTITPTSYNVISDSTGLHPDRMQWLTYKLTHMYFNCSAQVRVPAVCQYAHKLAFLAANSLHSQPHNSLTDTLYFL

Sup. Figure 2.

PCA (a) and MA (b) plot from the DESeq analysis published in Gimenez *et al.* , 2025 and re-used in this study

a

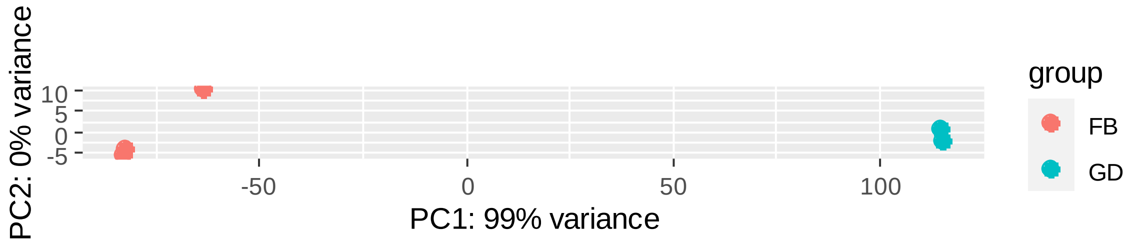


FB, fat body, GD, gonads of *Spodoptera frugiperda*

b

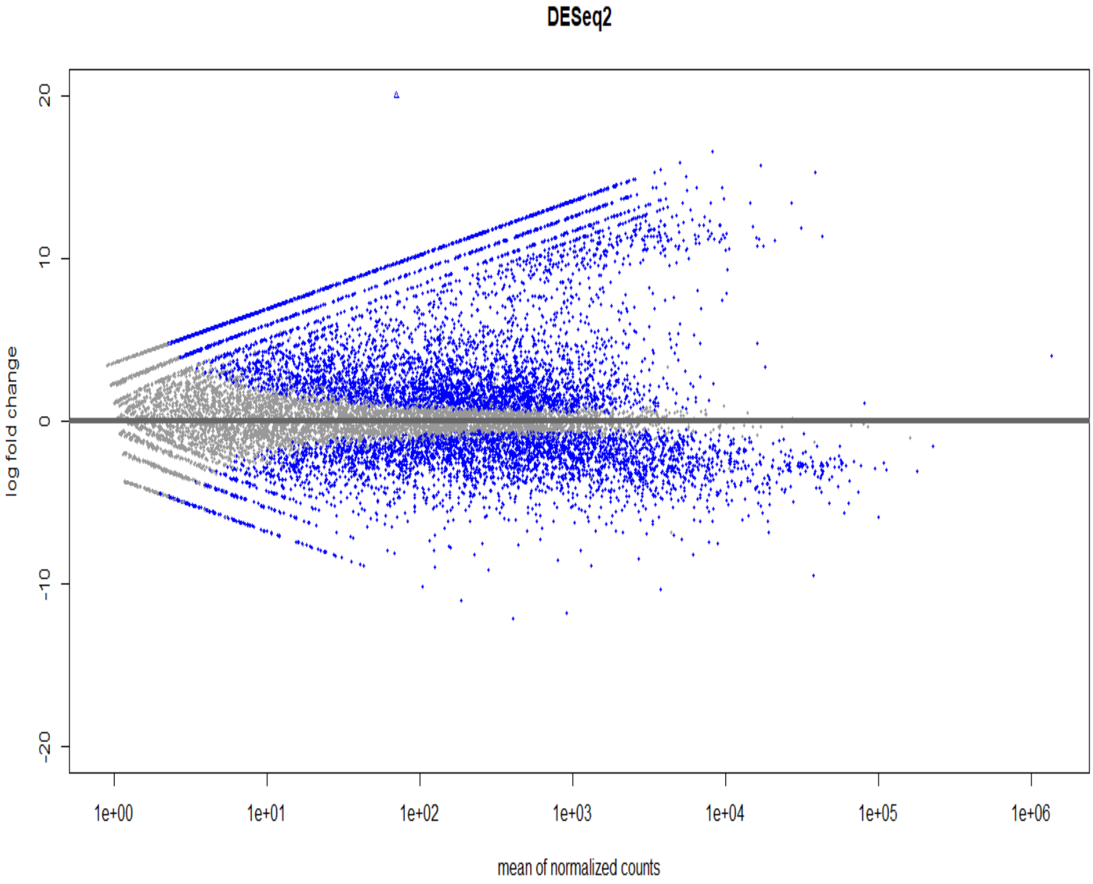


From Gimenez, S., Eychenne, M., Legeai, F., Gamble, S., and d'Alencon, E. (2025). Towards identification of a holocentromere marker in the lepidopteran model Spodoptera frugiperda. Chromosoma 134, 2. 10.1007/s00412-025-00828-2.

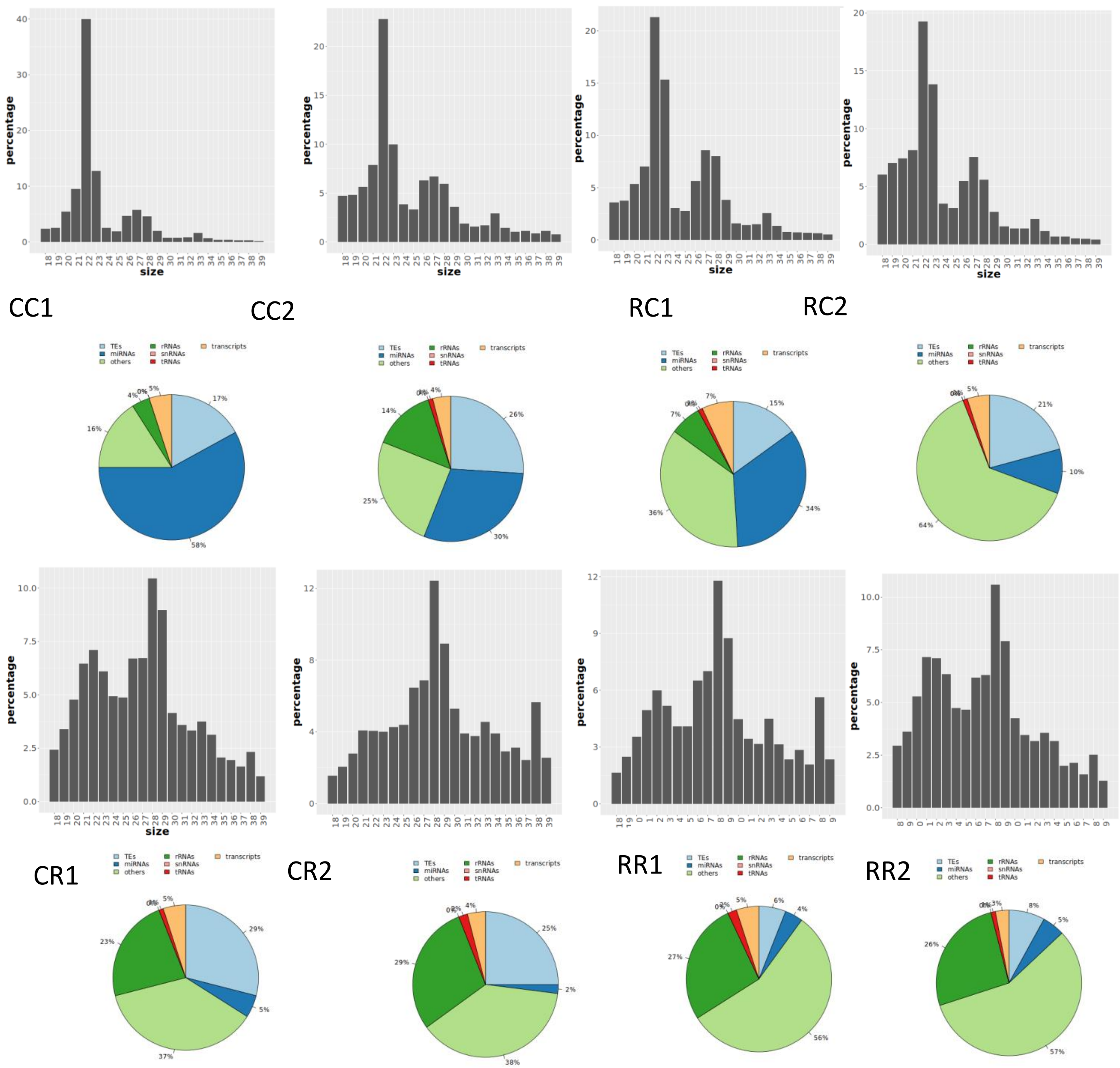


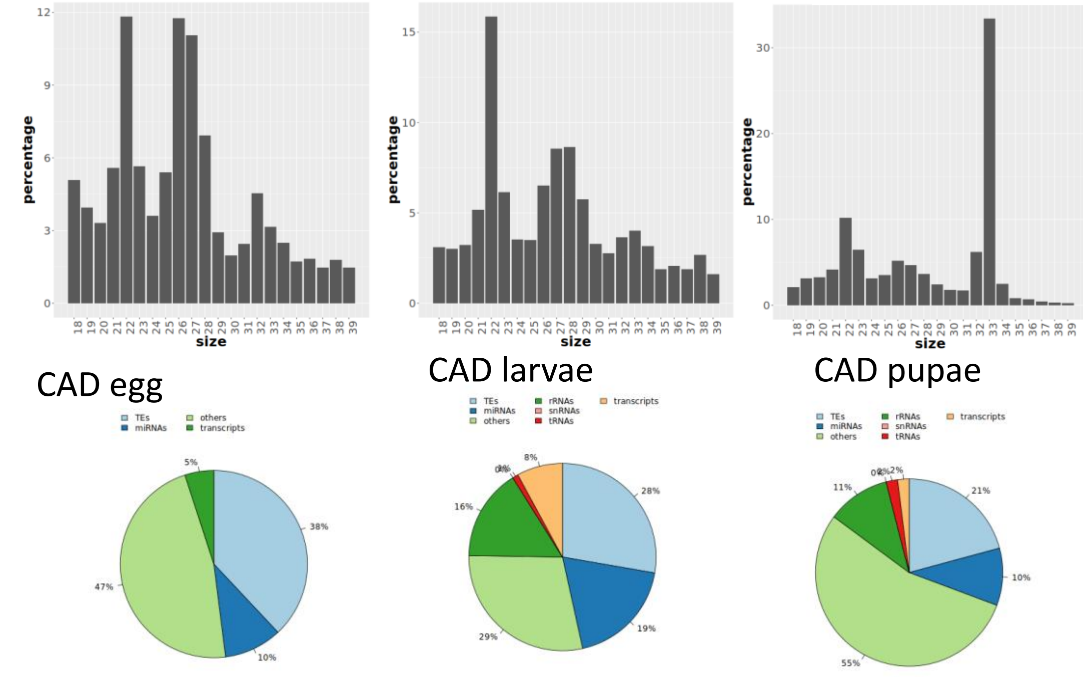


Sup. Fig. 3
Size profiling and pie-chart representing the proportion of sRNA sequences mapping to different genomic references

CC SfC on corn
CR SfC on rice
RC SfR on corn
RR SfR on rice
CAD SfC on artificial diet

Sup. Fig. 4. Long non coding RNAs mapping to TE and DNA repeats In SfC genome

| TE | Number of LNCRNA matching to TE |
|---|---|
| DXX | 9 |
| DTX | 67 |
| DHX | 53 |
| DMX | 2 |
| RXX | 39 |
| RLX | 33 |
| RYX | 5 |
| RIX | 51 |
| RSX | 12 |
| XXX | 7 |
| Potential hostgenes | 11 |
| NoCat | 31 |
| Total | 320 |

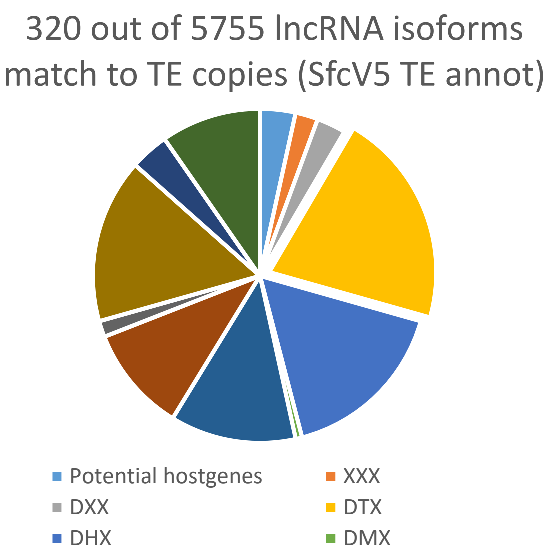


Retrotransposons classes in green: DIRS (RYX), LARD (RXX-LARD), LINE (RIX), LTR (RLX), PLE (RPX), SINE (RSX), TRIM (RXX-TRIM). DNA transposons classes in blue : Crypton (DYX), Helitron (DHX), MITE (DXX-MITE), Maverick (DMX), TIR (DTX). noCat (repeated sequence not classified at class AND order levels). XXX (sequence not classified at class level and with potential several orders)

### Sup. Fig. 5. GO term enrichment analysis of the coding genes regulated by piRNAs

#### A) Most specifically enriched terms

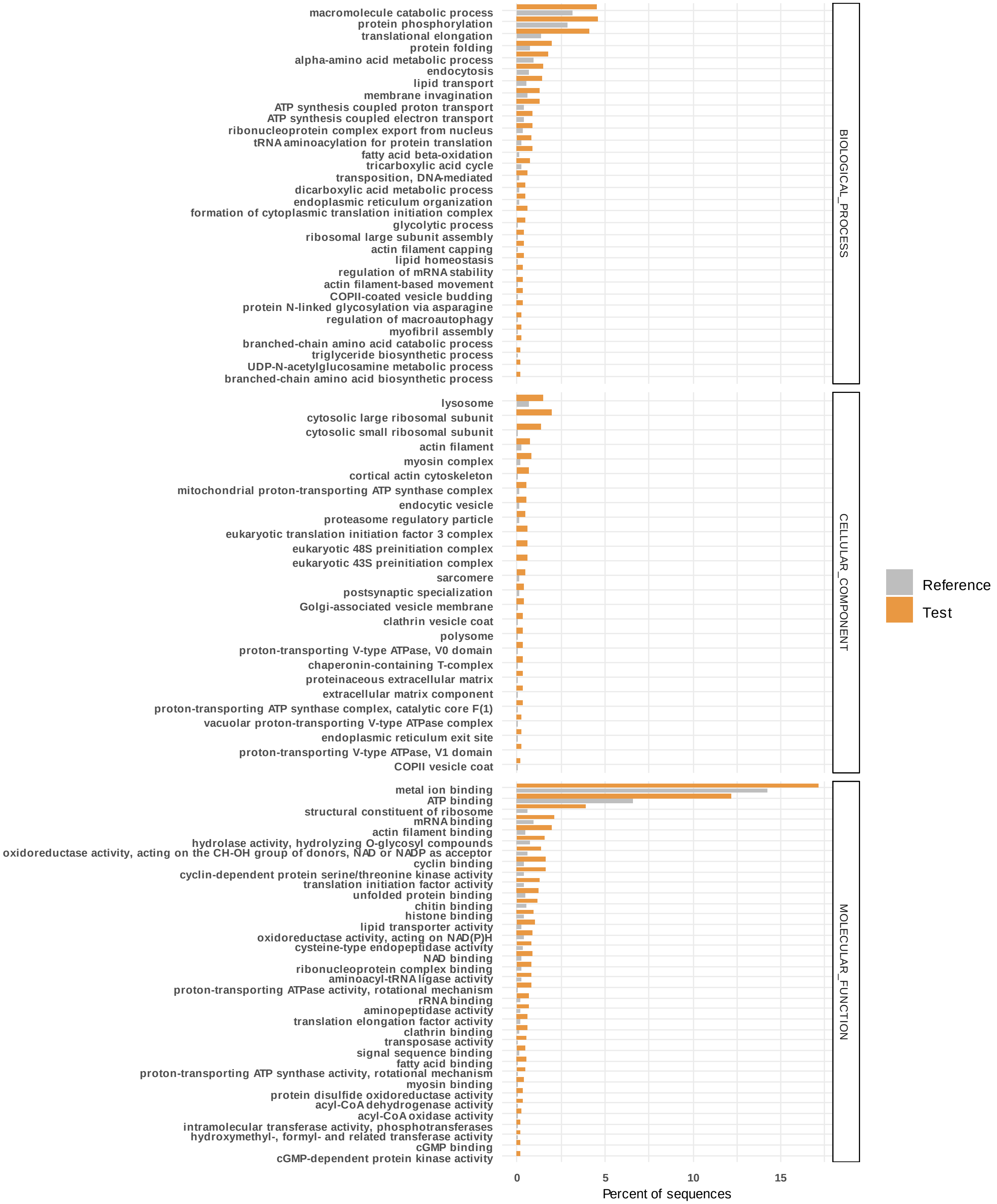

B) Functions of genes regulated by piRNAs

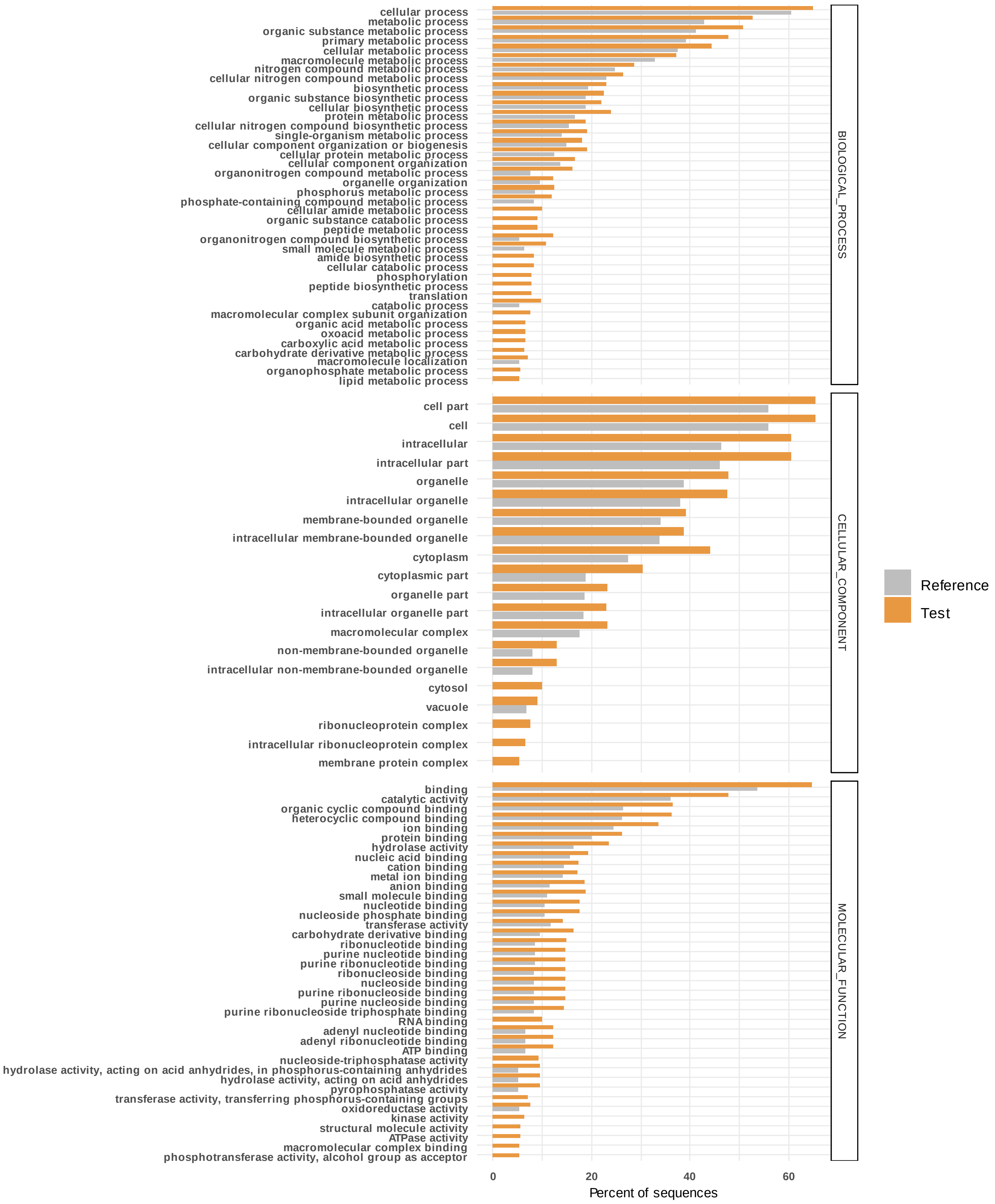
