## Supplementary table 5 for "A characterization of piARNs, their biogenesis and their targets in *Spodoptera frugiperda* (Lepidoptera, Noctuidae)"

| Genome assembly | SfC v5.0  (Nam Kiwoong, 2018) | SfR v3.0  (Nam et al., 2020) |
| --- | --- | --- |
| seqNb | 1000 | 1054 |
| totallen | 384358365 | 379902270 |
| N50 | 900335 | 1129192 |
| N90 | 196225 | 165330 |
| N95 | 108569 | 86988 |
| meanlen | 384358.365 | 360438.586 |
| medianlen | 160333 | 108017 |
| minlen | 8866 | 10636 |
| maxlen | 5279935 | 7849854 |
| %N | 0.0688 | 0.0006 |

Nam, K., Nhim, S., Robin, S., Bretaudeau, A., Nègre, N., and D'Alençon, E. (2020). Positive selection alone is sufficient for whole genome differentiation at the early stage of speciation process in the fall armyworm. Bmc Evolutionary Biology *20*.

Nam Kiwoong, N.S., Robin Stéphanie, Bretaudeau Anthony, Nègre Nicolas, d'Alençon Emmanuelle (2018). Divergent selection causes whole genome differentiation without physical linkage among the targets in Spodoptera frugiperda (Noctuidae). bioRxiv.
